# A study of phenotypic plasticity of *Saccharomyces cerevisiae* in natural grape juices shed light on allelic variation under balanced selection

**DOI:** 10.1101/288944

**Authors:** Emilien Peltier, Vikas Sharma, Maria Martí Raga, Miguel Roncoroni, Margaux Bernard, Vladimir Jiranek, Yves Gibon, Philippe Marullo

**Affiliations:** Univ. Bordeaux, ISVV, Unité de recherche OEnologie EA 4577, USC 1366 INRA, Bordeaux INP, Villenave d’Ornon, France; Biolaffort, Bordeaux, France; Universitat Rovira i Virgili, Departament de Bioquímica i Biotecnologia, Facultat d’Enologia de Tarragona, Tarragona, Spain; Wine Science Programme, University of Auckland, Private Bag 92019, Auckland, New Zealand; Department of Wine and Food Science, University of Adelaide, Urrbrae, South Australia 5064, Australia; INRA, University of Bordeaux, UMR 1332 Fruit Biology and Pathology, F-33883 Villenave d’Ornon, France

## Abstract

The ability of a genotype to produce different phenotypes according to its surrounding environment is known as phenotypic plasticity. Within different individuals of the same species, phenotypic plasticity can vary greatly. This contrasted response is due to allelic variations and is caused by gene-by-environment interactions (GxE). Finding the genes and the cellular functions that interact with the environment is a current challenge for better understanding the genetic bases of phenotypic plasticity. In order to study the impact of natural allelic variations having a contrasted but relevant effect in a changing environment, we investigated the phenotypic response of the wine yeast *Saccharomyces cerevisiae* fermented in various grape juices. In this study we implemented a QTL mapping program using two independent offspring (~100 progeny) in order to investigate the molecular basis of yeast phenotypic response in a wine fermentation context. Thanks to high throughput sequencing approaches, both populations were genotyped, providing saturated genetic maps of thousands of markers. Linkage analyses allowed the detection of 78 QTLs including 21 with significant interaction with the nature of the fermented juice or fermentation conditions. Molecular dissection of a major QTL showed that the sulfite pump Ssu1p has a pleiotropic effect and impacts the phenotypic plasticity of several traits. Both alleles have positive effect according to external condition in phenotypes related to yeast fitness suggesting an example of balanced selection. All together these results pave the way for exploiting and deciphering the genetic determinism of phenotypic plasticity.

## Introduction

Phenotypic plasticity, which is the ability of a genotype to produce distinct phenotypes in different environmental conditions, has been widely reviewed (West-Eberhard 1989; Pigluicci 2005; Nussey D. H., Wilson a. J., et Brommer J. E. 2007). It encompasses various aspects of the organism’s life including ontogeny (Sultan 2000), lifespan (Whitfield, Cziko, et Robinson 2003; Panowski et al. 2007), response to biotic (Agrawal 2001; Leal, Seehausen, et Matthews 2017) or abiotic (Shao et al. 2007; Nicotra et al. 2010) factors, pathogen or disease susceptibility (Shields et Harris 2000; Feinberg 2007; Roux, Gao, et Bergelson 2010), and animal behavior (Feinberg 2007; Sambandan et al. 2008). As a universal mechanism, phenotypic plasticity has been reported in humans (Feinberg 2007), animals (Dixon 1977; Spitze 1992; Panowski et al. 2007), plants (Sultan 2000), and fungi (Slepecky et Starmer 2009; Rai et al. 2017). The term of phenotypic plasticity can be used at different integrative levels. At the population level the phenotypic plasticity is the overall phenotypic response of a species to different environments. The genetic basis of those responses is mainly explained by transcriptional (Schlichting et Pigliucci 1993; Aubin-Horth et Renn 2009), post-transcriptional (Albertin et al. 2013), and/or epigenetic (Feinberg 2007; Rai et al. 2017) regulations. Phenotypic plasticity can also be considered at the individual level within a species (Pigluicci 2005; Nussey D. H., Wilson a. J., et Brommer J. E. 2007). In this case the phenotypic plasticity is measured for specific genotypes. The phenotypic response pattern observed is termed the reaction norm. Among individuals of the same species, non-parallel reaction norms are observed in animals (Veerkamp, Simm, et Oldham 1994; Sae-Lim et al. 2015), plants (Pigliucci et Kolodynska 2002; Rashwan 2016) and fungi (Marullo et al. 2009). These different patterns of response are due to the genotype-by-environment interaction (GxE) determined by allelic variations having different effects according to external conditions. Those variations entail individuals more or less adapted according to environment. Plasticity is therefore subject to the action of natural selection (Gotthard et Nylin 1995; Nussey D. H., Wilson a. J., et Brommer J. E. 2007). Understanding the evolution of plasticity is a complex issue because it depend on the relative frequency of different environments and their relative influence on overall fitness (Kawecki et Stearns 1993). However the genetic bases of GxE interactions can be investigated at a genomic level by QTL mapping programs carried out in various environmental conditions. This strategy has commonly been used for animals (Vieira et al. 2000; Gutteling et al. 2007), plants (Ungerer et al. 2003; Yang et al. 2016) and fungi (Smith et Kruglyak 2008; A. Bhatia et al. 2014; Aatish Bhatia et al. 2014; Wei et Zhang 2017). Although, many QTLs interacting with environment are often detected, the identification of the genetic basis of GxE at a gene level is far from being trivial and little molecular evidence has been reported in plants (Chiang et al. 2011; Campitelli, Des Marais, et Juenger 2016) and yeast (Gerke et al. 2010; Martí-Raga et al. 2017).

In agronomy, the concept of phenotypic plasticity, that affects the stability (robustness) of domesticated plant varieties and animal races across diverse uncontrollable macro environments, has been integrated a long time ago (Ceccarelli et al. 1994; Kang 1997; O’Neill, Swain, et Kadarmideen 2010; Sae-Lim et al. 2015). Apart from species of agronomic interest, fungi and in particular yeast are eukaryotic organisms with an important economic impact. The bakers’ yeast *Saccharomyces cerevisiae*, is by far the major industrial microorganism since it is involved in the production of numerous fermented foods including bread, wine and beer (Sicard et Legras 2011). These processes can be better controlled, by using industrial starters that have been subjected to genetic selection using breeding programs (Steensels et al. 2014). Depending on the composition of the fermentation matrix, yeasts are faced with various stresses and conditions inducing contrasted technological response. In winemaking, the initial composition of the grape must strongly impacts the alcoholic fermentation. Indeed, grape cultivars, enological practices, *terroir* and climate modulate biotic (populations size of various species including yeasts, molds, bacteria) and abiotic (sugar, nitrogen, lipid, vitamin concentration, oxygen, temperature, turbidity) factors and strongly shape the phenotypic variability of wine yeasts (Monk 1982; Cottrell et Lellan 1986; Gardner, Rodrigue, et Champagne 1993; Remize, Sablayrolles, et Dequin 2000; Fornairon-Bonnefond et al. 2003; Torija et al. 2003; Berthels et al. 2004; Luparia et al. 2004; Bell et Henschke 2005; Varela et al. 2012). Understanding how and why industrial starters have non-parallel norms of reaction is a critical challenge for industrial yeast selection. Therefore dissecting the phenotypic plasticity at the molecular level is of great importance for selecting individuals with favorable reaction norm able to ensure successful fermentation in a wide range of conditions. Recently the dissection of second fermentation kinetics QTLs offered the opportunity to find out allelic variations in two genes involved in pH homeostasis explaining why wine strains are differently adapted according to wine pH (Martí-Raga et al. 2017). In the present work, we applied a QTL mapping program aiming to identify QTLs interacting with environmental conditions. Eleven quantitative traits related to alcoholic fermentation were measured for two distinct populations of ~100 progeny each in three distinct conditions simulating diverse winemaking practices. The use of two independent backgrounds allowed estimation of the impact of parental divergence on QTL identification. High-density genetic maps generated by genome sequencing enabled QTLs to be detected at the gene level. Although most of the QTLs were robust to environmental changes, some striking GxE interactions were identified. One of these was explained at the gene level, revealing that allelic variations in the promoter of the gene *SSU1* are beneficial or not according to the fermentation conditions and provide an example of natural genetic variations under balanced selection among the wine strains group.

## Results

### Experimental design

The purpose of this study was to find out at a large scale QTLs interacting with the environment by using the model yeast *S. cerevisiae*. As QTLs can be readily used in yeast breeding for strain improvement, this work was carried out in an enological context by measuring the phenotypic plasticity of wine related strains in three contrasted conditions met in winemaking. Two particular effects were investigated: (i) the phenotypic plasticity of yeast fermenting red or white grape musts (GM) was estimated by comparing M15_Sk (red) *vs* SB14_Sk (white) conditions, reflecting common types of grape juice fermented around the world. (ii) the phenotypic response to micro-oxygenation (µ-Ox) in accordance with enological practices (Peltier et al. 2018) was estimated by comparing M15_Sk (shaken) *vs* M15 (unshaken). In order to have the broadest landscape of phenotypic plasticity, we performed QTLs mapping in two distinct genetic backgrounds derived from commercial starters widely used in wine industry (SB, GN, M2 and F15). The full experimental design is summarized in Fig 1. Two main questions were then addressed. First, we evaluated if the genetic and/or phenotypic characteristics of parental strains impact the architecture of quantitative trait determinism. Second, we investigated if and how yeast strains display different norms of reaction according to the grape juice and the fermentation conditions used.

**Fig 1.**
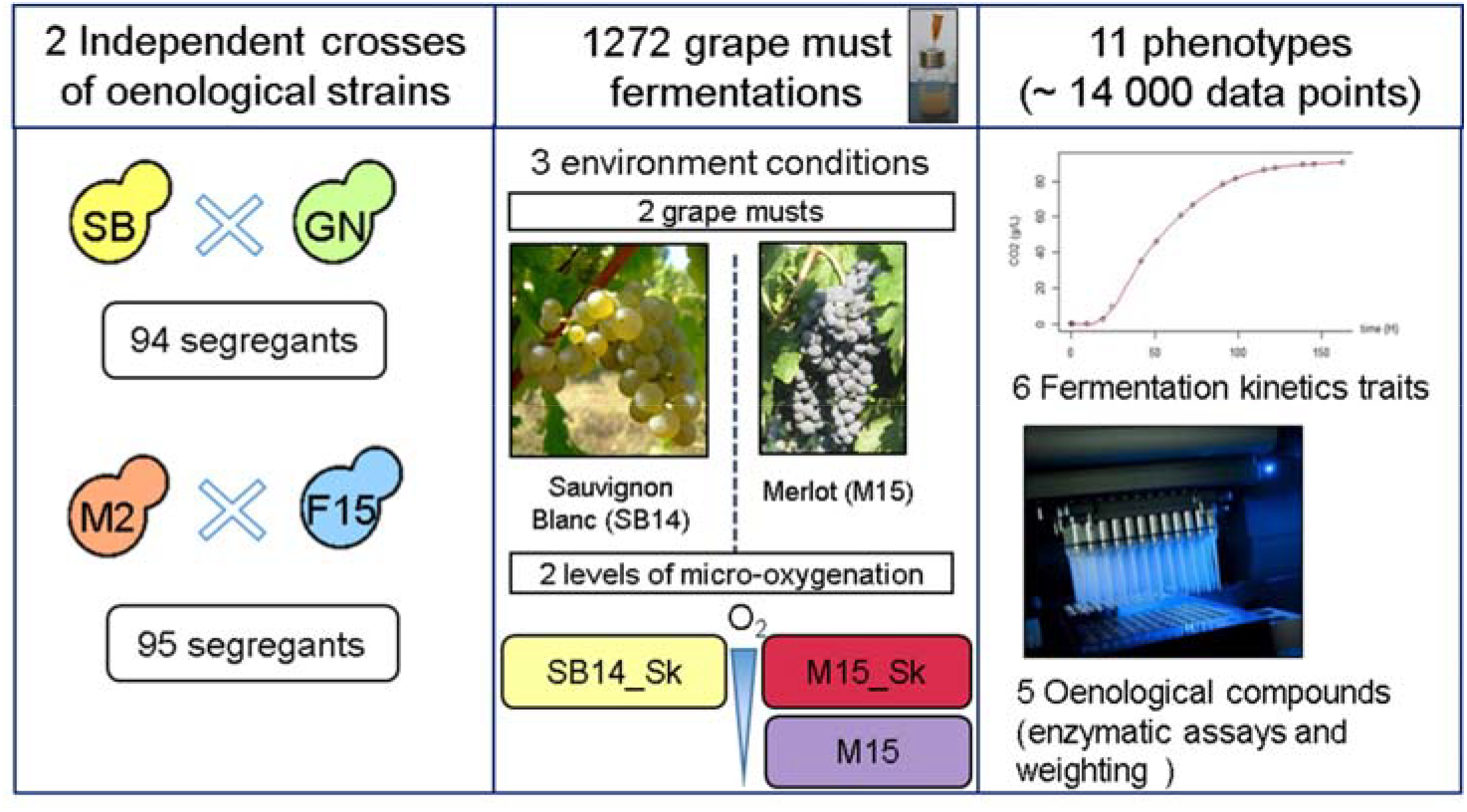
Experimental design. Four diploid homozygous strains (SB, GN, F15, M2) were used to generate two independent hybrids (SBxGN and M2xF15) and their 94 and 95 segregants, respectively. The whole genome sequence of each parental strain and each progeny was obtained by high throughput sequencing and analyzed to get saturated genetic maps (3433 and 8378 markers, respectively). The phenotypic characterization of the strains was carried out in three conditions (SB14_Sk, M15_SK and M15) in small volume vessels. Fermentation time course was monitored and six traits were computed: lag phase (*Ip*), time required to produce 35, 50 and 80 g.L^−1^ of CO_2_ (*t35g, t50g* and *t80g*, respectively) and glucose consumption rate in the first and the second part of the fermentation (*V15_50* and *V50_80*, respectively). The concentrations of five end-point metabolites were also assayed by enzymatic methods (*acetic acid, glycerol, pyruvate* and *SO*_*2*_) or by weighting (*CO*_*2*_*max*). Grape pictures were reprinted from CC-BY licences, the copyright holders being Lebowskyclone (Merlot) and User:Vl (Sauvignon Blanc).

### Genetic and phenotypic relationships of the parental strains

In order to estimate the genetic relationships between the four parental strains, 15 polymorphic microsatellite markers were used (Legras et al. 2007). The codominant allele set of each parental strain was compared to those of 96 Commercial Wine Starters (CWS) encompassing the overall diversity of the *S. cerevisiae* wine group (S1 Table). This dataset was used to build a tree using Bruvo’s distance (Fig 2, panel A). The pairwise Bruvo’s genetic distance between all the strains ranged from 0 (identical strains) to 0.924, with an average of 0.552. According to previous studies, the wine strain population is poorly structured. Four groups of strains (nodes with bootstrap values higher than 85%) were found. One of these groups corresponds to the Champagne-like strains, which have specific genetics features (Legras et al. 2007; Borneman et al. 2011). A second encompasses the strain Actiflore Bo213 and the parental strain SB. As expected, the parental monosporic clones (SB, GN, M2, F15) are closely related to their respective commercial ancestors (Actiflore BO213, Zymaflore VL1, Enoferm M2 and Zymaflore F15, respectively). Considering the overall genetic diversity of the *S. cerevisiae* species, the wine yeast cluster constitutes quite a homogenous group (Legras et al. 2007; Borneman et al. 2011). However, within this group the genetic relationships between parental pairs are different. The strains GN and SB are more divergent than the strains M2 and F15 (Bruvo’s genetic distance = 0.611 and 0.216, respectively; Fig 2, panel B). Similar results were obtained by calculating the % identity of a subset of 5281 polymorphic SNPs extracted from parental genomes (Sharma, personal communication). The relative phenotypic distance between parental strains was estimated using the phenotypic values of eight fermentation traits measured in five grape musts for 35 strains previously obtained (Peltier et al. 2018). In order to represent the genetic diversity of commercial starters, the sampled strains belong to the different branches of the tree. The pairwise Euclidian’s phenotypic distance between all the strains ranges from 0.73 (similar strains) to 9, with an average of 3.7. The strains SB and GN are phenotypically very different (9) while F15 and M2 have more similar phenotypes (3). All together, these results demonstrate that the hybrids used for QTL mapping (M2xF15 and SBxGN) were obtained from independent strains showing either moderate or high genetic and phenotypic distances within the wine strain group.

**Fig 2.**
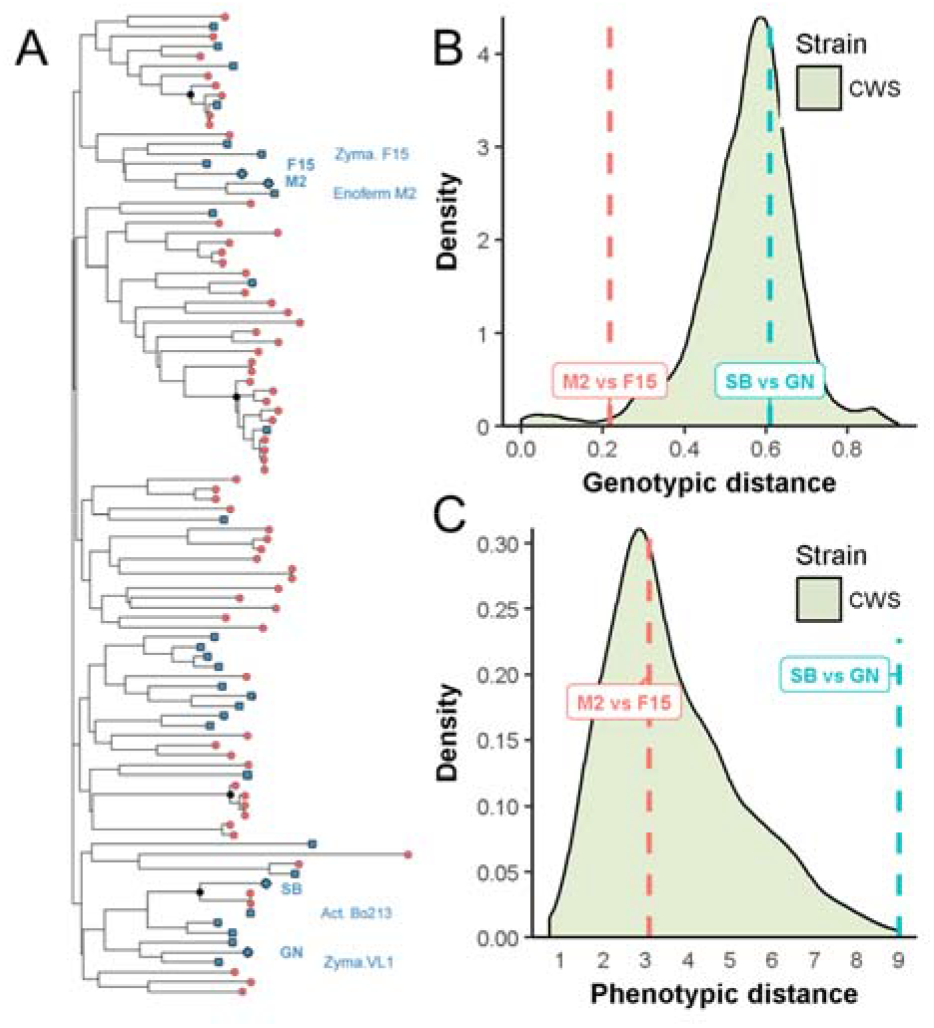
Genetic distance between parental strains. Panel A. Genetic relationships between 96 commercial wine yeast strains and the four parental monosporic clones GN, M2, F15 and SB used in this work (diamonds). The dendrogram tree was built using Bruvo’s distance and Neighbor-Joining’s clustering strains from the inheritance of 15 polymorphic microsatellites described by Legras et al [55]. Black dots represents nodes encompassing group of strains highly similar (bootstrap values >85). Among the strains genotyped, 31 Commercial Wine Starter (CWS) have been previously phenotyped and covered all the branches of the tree (blue squares). Panel B. The distribution of Bruvo’s distance for all genotyped strains. The relative distances between SB vs GN and M2 vs F15 are 0.61 and 0.22, respectively. Panel C. The distribution of the phenotypic distance computed for 35 strains (31 CWS and the four parental strains). The data used eight traits in five grape musts and were obtained from (Peltier et al. 2018). Dashed lines shows the relative distance between parental pairs SB vs GN and M2 vs F15.

### Meiotic recombination emphasizes phenotypic novelty in either tested cross

By implementing alcoholic fermentations in a small volume, eleven heritable traits were measured in three grape musts/conditions for 195 strains in duplicate constituting a data set of 14,000 data points (S2 Table). From fermentation kinetics, six representative traits were extracted: lag phase duration (*lp*), time to produce various amount of CO_2_ (*t35g*, *t50g* and *t80g*) and fermentation rate during the first (*V15_50*) and the second part of the fermentation (*V50_80*). Moreover four metabolites present at the end of the fermentation were measured by enzymatic assays: *acetic acid, glycerol, pyruvate* and *SO*_*2*_. All fermentations were completed since the residual sugars were less than 2 g.L^−1^. The progress of alcoholic fermentation was measured by estimating the *CO*_*2*_*max* produced, which is stoichiometric to ethanol. The inheritance of the eleven quantitative traits was calculated for each cross and condition (Fig 3, panel A). Most of the traits were highly heritable since 75 % of them had h^2^ values higher than 0.5. Considering all the traits, each cross-by-environment combination had a distinct heritability profile and hierarchical clustering grouped them by environment rather than by cross (Fig 3, panel A).

**Fig 3.**
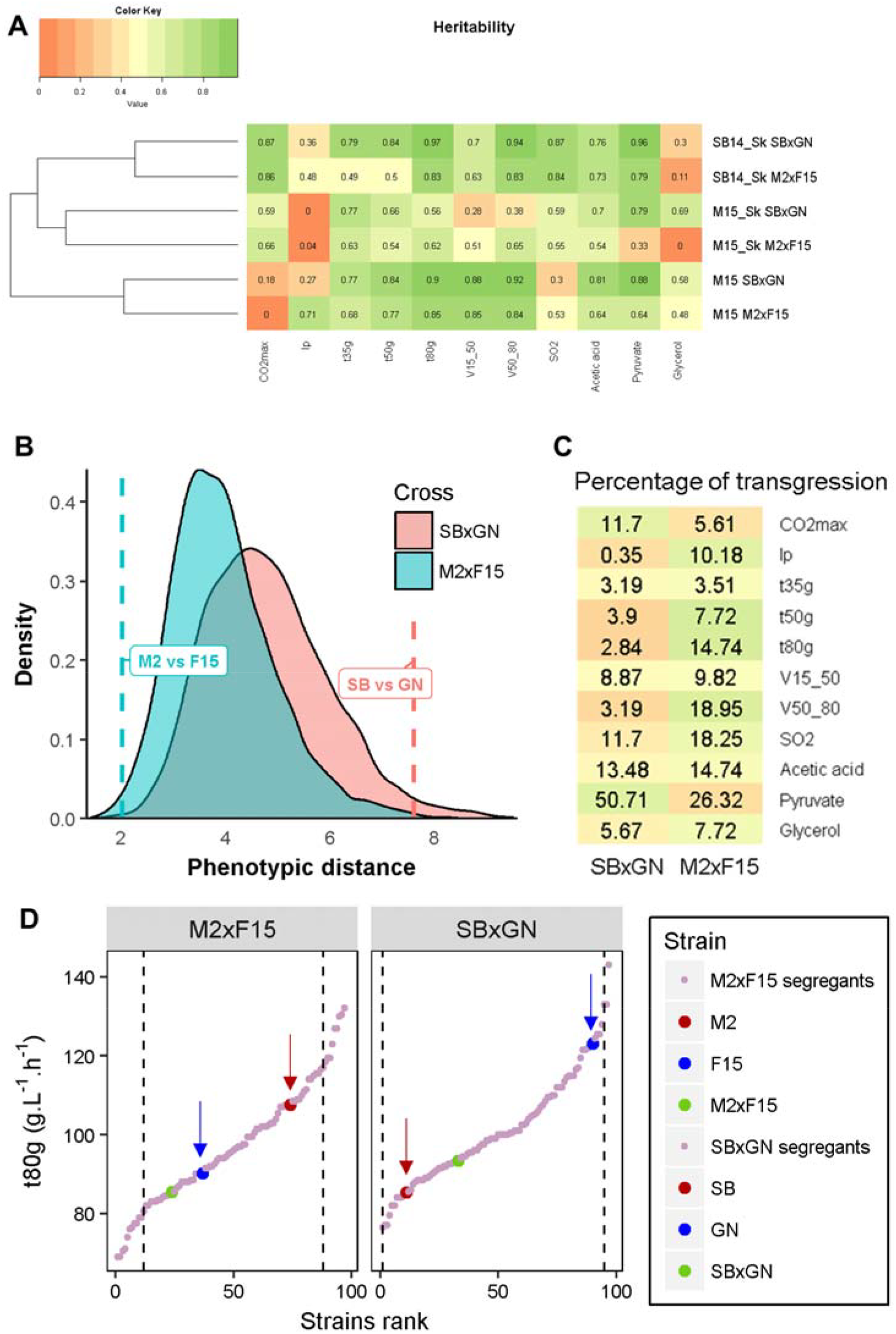
Phenotypic distribution patterns in the two progeny populations. Panel A. Heritability calculated for each trait according to cross and condition. Panel B. Dashed lines show the phenotypic distance between parental strains. Shaded areas show the distribution of the phenotypic distance within progenies. Panel C. Average percentage of transgression per trait in the three conditions according to the cross. Color key is scaled by trait, dark green means high transgression level and dark red low transgression level. Panel D. The distributions of *V50_80* (M15_Sk condition) illustrate the transgression discrepancy observed between the two crosses. Each dot represents a strain value ordered according to the rank. Parental phenotypes are indicated by arrows. Dashed vertical lines represent the upper and lower limits set for considering transgressive progeny clones.

The overall level of phenotypic segregation of both sets of offspring was estimated by computing the Euclidean distances within each progeny clone. The trait values for each condition were normalized (center-reduced) for preventing scale effects and only one kinetics trait (*t80g*) was used, since most of them were strongly correlated (see method) (Fig 3, panel B). The parental pairwise distances confirmed that SB vs GN are phenotypically more divergent than M2 vs F15 (Euclidian distance = 7.6 and 2, respectively). These contrasted relationships do not impair a wide segregation in both progeny populations. Indeed, the distributions of phenotypic distances are similar between the M2xF15 and SBxGN progenies illustrating that meiotic segregation generated a burst of phenotypic novelty (Fig 3, panel B). Surprisingly, the phenotypic magnitude generated was higher than those observed for the panel of 31 representative commercial starters phenotyped in the same conditions (S1 Fig). As a direct consequence, the percentage of transgressive progenies was much higher in the M2xF15 background, except for *CO*_2_*max* and *pyruvate* (Fig 3, panel C). The percentage of transgression in both offspring is illustrated for the *t80g* measured in M15_Sk in Fig 3, panel D. Overall, these biometric analyses indicate that meiotic segregation generated an important and similar phenotypic diversity despite the contrasted relationships of the parental pairs. In contrast, the relative phenotypic distance of parental strains had a strong effect on the transgression level observed. In both cases, the phenotypic variability that was generated significantly exceeded that found for a wide panel of commercial starters, thus underlining the power of meiotic segregation to generate phenotypic novelty.

### Distribution of individual reaction norms in segregating populations

The phenotypic plasticity was then analyzed for both progeny populations. First, the impact of the cross was tested using the linear model LM1 that estimates the effects of *cross* (C) and *environment* (E) as well as their primary interaction (C*E) (S3 Table). Analysis of variance shows that the sum of C and C*E had very low effects (below 3.4 % in average) for all phenotypes. This result indicates that both crosses have the same phenotypic response to environmental changes. The environmental conditions (μ-Ox and GM) affected the traits in a different manner. Three representative patterns of phenotypic response are shown in Fig 4; the full details are given in Table S4. As the two grape musts (SB15 and M14) used have different a composition in terms of SO_2_ (34 mg.L^−1^ *vs* 46 mg.L^−1^) and initial sugars (194 g.L^−1^ *vs* 219 g.L^−1^), the production of SO_2_ and CO_2_ (Fig 4, panel A) was directly impacted by the grape juice matrix. In contrast, other traits such as *glycerol* (Fig 4, panel B) and *acetic acid* were mainly impacted by μ-Ox. Indeed, in accordance with previous reports, an increased μ-Ox level increases glycerol production and decreases acetic acid production (Gardner, Rodrigue, et Champagne 1993; Remize, Sablayrolles, et Dequin 2000; Valero et al. 2002; Peltier et al. 2018). Some traits were influenced by both parameters (GM and μ-Ox), including kinetic parameters like *V50_80* (Fig 4, panel C). The influence of sugar content and μ-Ox on kinetic parameters has also been reported previously (Fornairon-Bonnefond et al. 2003; Varela et al. 2012). Finally, *pyruvate* production was only slightly impacted by the environmental conditions.

**Fig 4.**
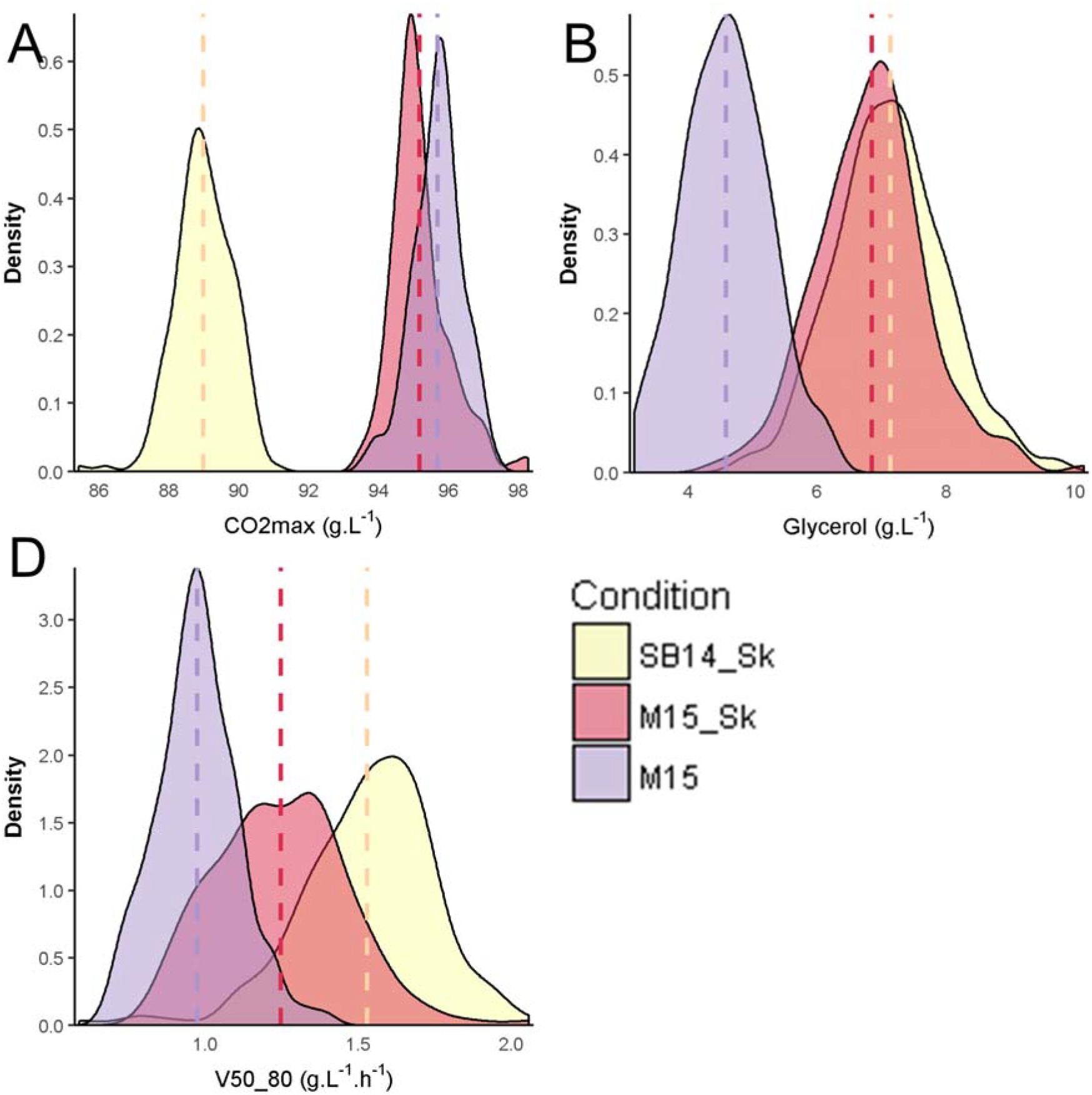
Environnemental impact on quantitative traits. Distribution of the phenotypic values according to fermentation conditions. Vertical dashed lines indicate the average value in each condition. *CO2max* is mainly impacted by grape must (Panel A), *glycerol* mainly by micro-oxygenation (Panel B) and *V50_80* by both (Panel C).

In order to estimate the reaction norm at a strain level, the variation of each trait was decomposed by a second linear model (LM2 see methods). In this model, the overall genetic impact was decomposed into S and S*E components. S represents the constant strain effect across the three conditions (*i.e.* parallel norm of reaction) whereas S*E represents the phenotypic plasticity among strains (*i.e.* non-parallel norm of reaction). The analysis of variance estimated the effect of strain (S) and *environment* (E) factors as well as the *strain x environment* (S*E) interaction (Table 1). The model explained a great part of phenotypic variation (between 62.8 and 96.5 % according to the trait). For the majority of traits, environment accounted for most of the variation. However, the constant strain effect S (parallel norm of reaction) also explained an important part of phenotypic variability (up to 52.4 % for *pyruvate*). For some traits, an important phenotypic plasticity was observed. This is the case for *lp* (26.1 %) and *acetic acid* (20.6 %), for which a non-parallel norm of reaction (S*E) made a non-negligible contribution. Interestingly, the genetic weight for CO_2_ production kinetics increased with the fermentation ongoing. Indeed, the sum of S + S*E explained 17.1 % of the variance for t35g and linearly increased to 37.6 % for *t80g*.

**Table 1.** Analysis of variance of the model LM2 for the 11 phenotypes with 189 strains and three conditions of fermentation. Percentage of variance explained by the LM2 model. Signifiance codes: p. val < 0.001 = ***, p. val < 0.01 = **, p. val < 0.05 = *.

|  | CO<sub>2</sub>max | Lp | t35g | t50g | t80g | V15_50 | V50_80 | SO<sub>2</sub> | Acetic acid | Pyruvate | Glycerol |
| --- | --- | --- | --- | --- | --- | --- | --- | --- | --- | --- | --- |
| E | 90.6 *** | 12.8 *** | 77.2 *** | 71.2 *** | 53.3 *** | 59.4 *** | 53.3 *** | 54 *** | 22.1 *** | 9.2 *** | 55.3 *** |
| S | 2.7 *** | 23.9 *** | 10.8 *** | 13.9 *** | 21.4 *** | 17.8 *** | 19.6 *** | 18.3 *** | 36.1 *** | 52.4 *** | 13.6 *** |
| S*E | 3.2 *** | 26.1 | 6.3 *** | 8.4 *** | 16.2 *** | 11.7 *** | 17.8 *** | 12.6 *** | 20.6 *** | 17.9 ** | 13.2 |
| Residuals | 3.5 | 37.2 | 5.6 | 6.5 | 9.2 | 11.2 | 9.4 | 15.1 | 21.2 | 20.5 | 17.9 |

In order to better determine the parallel (S) and non-parallel (S*E) norms of reactions, the phenotypic plasticity of the 189 strains was organized by k-means clustering on the basis of a matrix of correlation distance (see methods). This procedure clustered each progeny clone according to its phenotypic response against environment. This procedure clustered each progeny clone according to the similarity of their phenotypic response. Divergent norm of reaction patterns were identified for each trait (S2 Fig). For *acetic acid*, four clusters were obtained, they mainly contained strains with plastic response to micro-oxygenation (μ-Ox: 101 strains), to grape juice (GM: 44 strains), and to the combined effect of GM and μ-Ox (40 strains) (Fig 5). Strikingly, only seven strains (3.7%) appeared robust regarding environmental conditions (Robust cluster). In the four clusters, a similar number of progeny clones was found for the two crosses (chi-squared test, α = 0.05) indicating that the plastic response pattern is not cross specific in this experiment. From an enological point of view, the magnitude observed is quite relevant since a 0.1 g.L^−1^ acetic acid difference may impact wine quality (Ribéreau-Gayon et al. 2006). For example, micro-oxygenation had a positive impact for the strains of the cluster μ-Ox and the cluster GM + μ-Ox by significantly decreasing *acetic acid* production (Kruskal-Wallis rank sum test, p val < 0.05). In contrast the strains of cluster GM should be preferred for fermenting white grape juice rather than red matrices (Kruskal-Wallis rank sum test, p val < 0.05).

**Fig 5.**
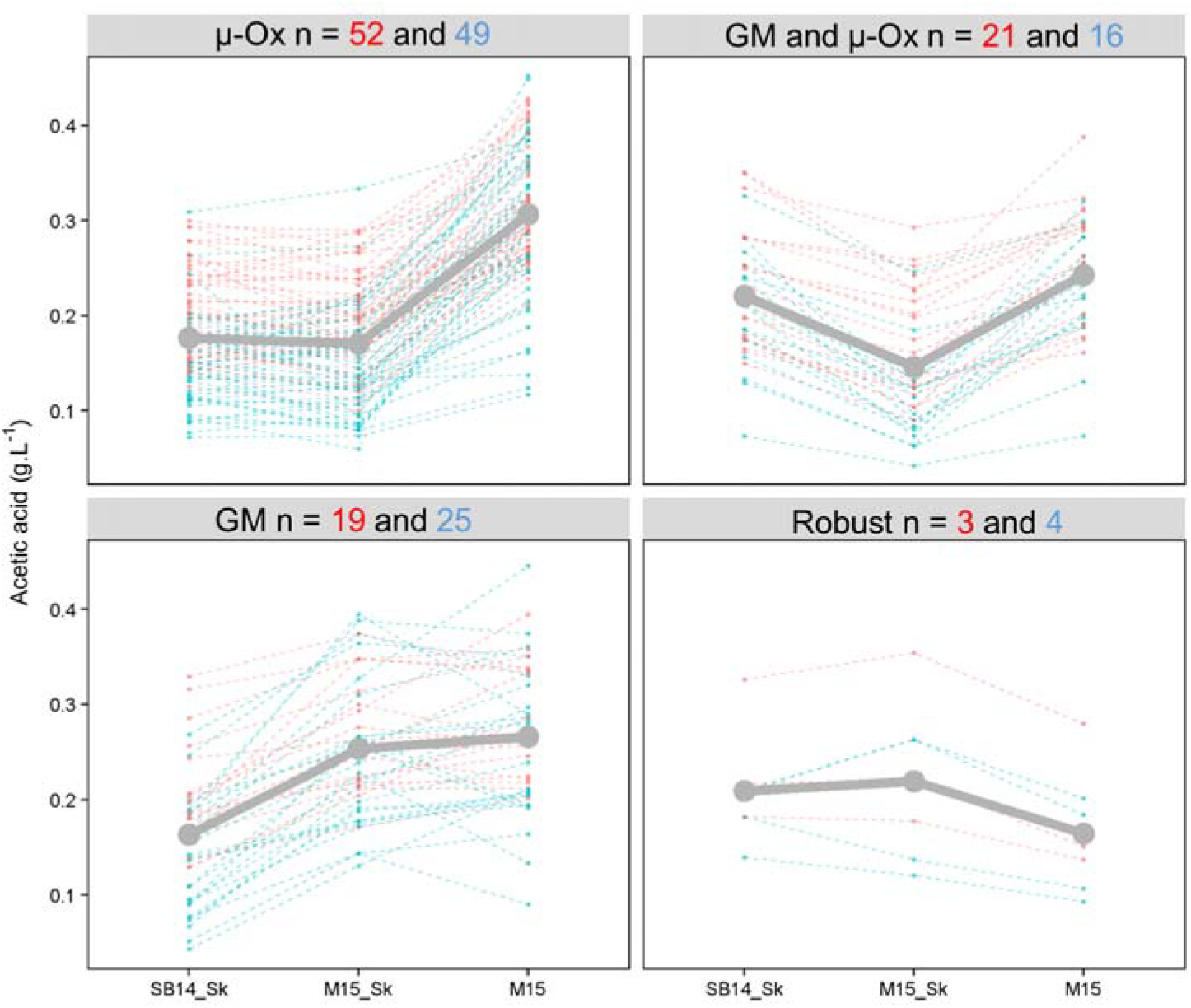
Divergent norm of reaction for acetic acid. Norm of reaction of each individual is shown by dotted line and cluster. They are colored according to the cross (blue for SBxGN offspring and red for M2xF15 offspring). A grey solid line shows the average norm of reaction of each cluster. Number of strains within each cluster is indicated and colored according to the offspring (red for M2xF15 offspring and blue for SBxGN offspring).

### Genetic architecture of complex traits is explained by a similar number of QTLs

The genetic determinism of the eleven traits was elucidated by QTL mapping in both crosses. Phenotypes were linked to segregating genetic markers identified by whole genome sequencing ((Roncoroni 2014) and this work). The high marker density (0.3 and 0.6 markers/kb for M2xF15 and SBxGN, respectively) ensure a precise localization of QTLs (Bloom et al. 2013; Wilkening et al. 2014; Martí-Raga et al. 2017). Interval mapping was carried out by applying a Haley-Knott regression model. This model estimates the effect of each *QTL* detected, the effect of each *Environment* (SB14_Sk, M15, M15-Sk) and the interaction effect between *QTL* and *Environment* (Fig 6, panel A). Statistical thresholds (FDR = 5%) were estimated by 1000 permutation tests (Churchill et Doerge 1994). Since the fermentation kinetics traits were partially correlated (*t35g, t50g, t80g, V15_50* and *V50_80*), we found numerous QTLs corresponding to closely related markers. In such cases, a unique QTL was considered in a window of 10 kb and was assigned to the kinetic trait showing the lowest p value (see methods). For ease of discussion, all the QTLs found for fermentation kinetics traits were then grouped into the “Kinetics” category. We mapped 78 unique QTLs in the two offspring (S5 Table). The efficiency of the multi-environment model was compared to the simplest models, in which only one environmental condition was used. The multi-environment model strongly increased detection power since 45 additional QTLs were detected by this method (Fig 6, panel B). The number of QTLs detected ranged from three (for *CO*_*2*_*max*) to 28 (for *Kinetics*) illustrating the complex genetic determinism of the traits investigated and was correlated with the trait heritability (S3 Fig).

**Fig 6.**
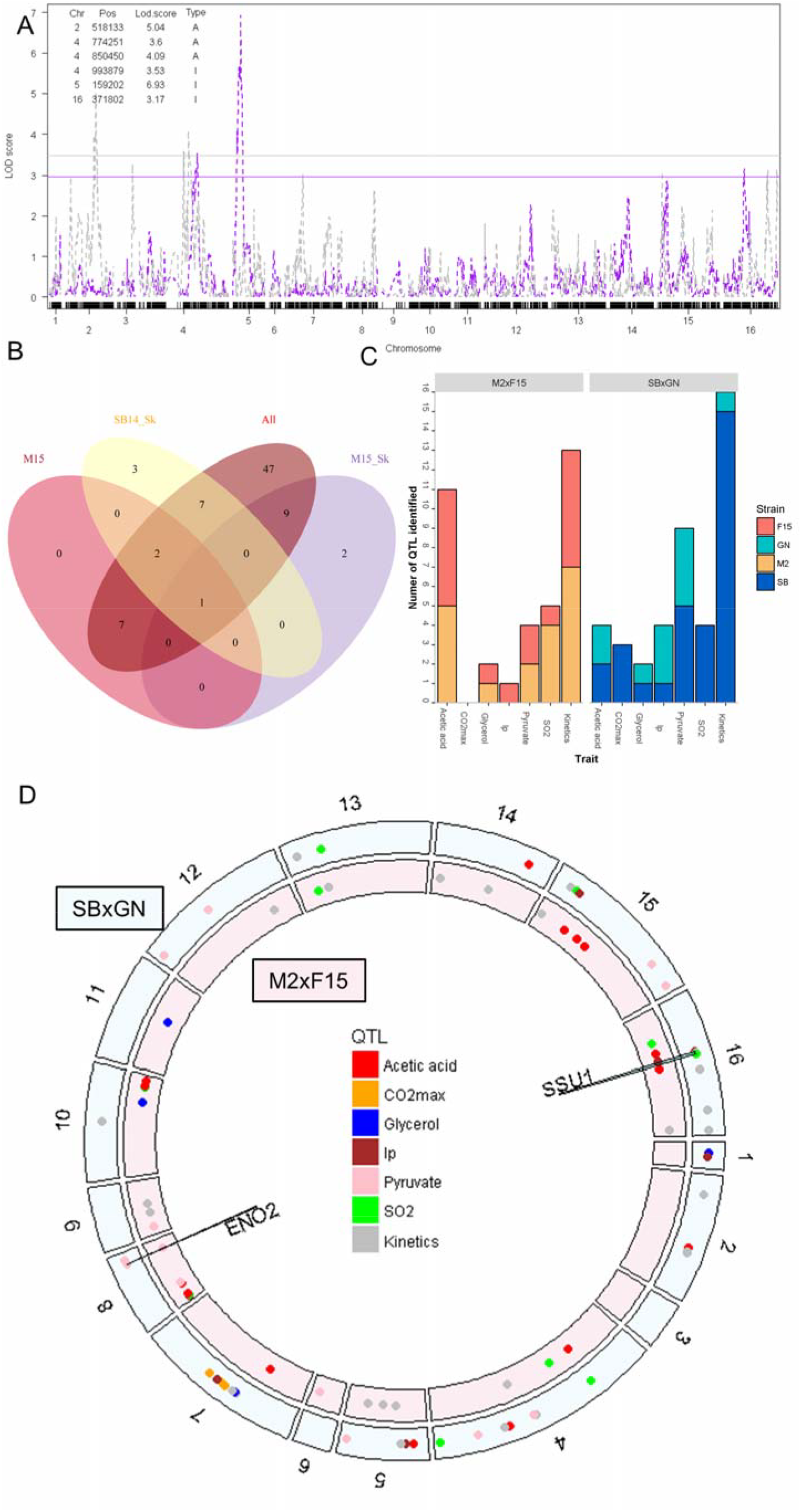
Number of QTLs identified according to cross or environmental conditions. Panel A. LOD score obtained with SBxGN offspring for *V50_80*. In grey, LOD score for the model with environment as additive variable, in pink, LOD score for the model with environment as interactive variable. The corresponding horizontal colored lines represent the 5% FDR threshold. The table details the QTLs identified by indicating their chromosome (Chr), the position of the marker with the highest LOD score within the QTL peak (Pos), the maximum LOD score within the QTL peak (Lod.score) and the model that detected the QTL (Type, A = additive, I = interactive). Panel B. The Venn diagram presents the number of QTLs identified according to the model used. The *All* ellipse corresponds to the multi-environmental model while M15, SB14_Sk and M15_Sk ellipses counts QTLs detected only using a single condition. Panel C. The bar chart presents QTLs identified according to cross and trait. Common QTLs identified for *t35g, t50g, t80g, V15_50* and *V50_80* are pooled in Kinetics category and are counted only as one QTL. QTLs are colored according to the parental strains that possess the favorable allele in an enological context. Panel D. Distribution of the QTLs identified along the genome according to the cross (M2xF15 inner track, SBxGN outer track). Each point indicates a QTL and is colored according to the trait. The two QTLs that colocalize in the two crosses and linked to the *SSU1* and *ENO2* genes are indicated.

Despite the contrasted F1-hybrids used (Figs 2 and 3), a similar number of QTLs was mapped in both progeny populations, with 36 and 42 QTLs for M2xF15 and SBxGN, respectively (Fig 6, panel C). The positive contribution of each parental strain was evaluated considering the suitable trait value expected according to enological practices. The numbers of positive alleles inherited from M2, F15, SB and GN was 19, 17, 31 and 11, respectively. According to the trait and the cross, the positive alleles were inherited from both or mostly one parent. A noteworthy unbalance was found for kinetics traits since 15/16 of the positive alleles were inherited from the fastest parental strain (SB) in the SBxGN cross whereas only 6/13 were inherited from the fastest parental stain (F15) in the M2xF15 cross (Fig 6, panel C). In order to find QTLs common to both progeny populations, we sought QTLs impacting the same trait in a 20-kb window (Fig 6, panel D). Only two QTLs were shared by the two progeny populations. The first locus was detected for *SO*_*2*_, *lp* and *V50_80* and is closely linked to the gene *SSU1* (markers: XVI__SBxGN__371802 and XVI__M2xF15__355235). In the SBxGN progeny, this QTL peak was strongly linked with the marker XV__SBXGN__172951 due to a reciprocal translocation event previously described for impacting the lag phase (Zimmer et al. 2014). The linkage between *SSU1* and the SO_2_ content at the end of the alcoholic fermentation is consistent with its molecular function. Indeed this gene encodes a sulfite transporter (Ssu1p) able to pump out the SO_2_ accumulated in the cytoplasm (Park et Bakalinsky 2000). The GN inheritance of this QTL, which reduces lag phase by increasing the *SSU1* expression (Zimmer et al. 2014) also increases the final SO_2_ concentration (Fig 6). Interestingly, in the M2xF15 hybrid the marker XVI__M2xF15__355235 was strongly linked to the marker VIII__M2xF15__6499. These two loci correspond to a translocation event (VIII-t-XVI) involving the gene *SSU1* previously described by Perez Ortin et al. (Pérez-Ortín et al. 2002) and present in M2xF15 hybrid (Roncoroni 2014).

The second common QTL mapped concerns the production of pyruvate and is located on chromosome VIII. This QTL explains 19.6 and 30 % of the total variance of pyruvate production in M2xF15 and SBxGN offspring, respectively. Interestingly, the markers VIII__SBxGN__446336 and VIII__M2xF15__449469 were located within 5 kb of the gene *ENO2* which is directly involved in pyruvate biosynthesis. Until now, the sequence analysis of the four parental strains did not reveal any relevant candidate SNP close to this gene. All together, these results suggest that most of the mapped QTLs are background dependent. Moreover, the similar number of QTLs in both progenies demonstrates that the mapping efficiency is no related, neither to the genetic, nor to the phenotypic distances between parental strains. However, a balanced contribution is more frequent when the strains are phenotypically similar.

### QTLxEnvironment interactions shape the phenotypic variability

The genetic determinism of phenotypic plasticity was then investigated at a genomic scale using the linear model LM3 (see method). For each QTL mapped, the constant genetic effect across environment (G), the interaction effect with grape must (GM), as well as the interaction effect with micro-oxygenation (μ-Ox), were estimated by analysis of variance. From this analysis, QTLs could be sorted as robust (no interaction) or interactive (significant interactions with GM and/or μ-Ox). The overall *GxE* pattern was shown for both crosses (Fig 7). The less the QTLs interacted with GM and μ-Ox, the closer they were from the bottom left corner of the ternary plot. Most of the QTLs (57/78) were quite robust to environmental changes (S6 Table). This was the case for the QTL V__M2F15__311505 that is one of the most robust. Indeed, progeny clones with both M2 and F15 inheritance showed parallel norms of reaction for *V50_80*. In contrast, 21 *GxE* interactions were detected; most of them (17) are due to the grape must composition. These interactions are mostly a “scale effect” as shown for the inheritance of the marker XVI__SBxGN__879639, which had an impact on *t35g* only in unshaken conditions (Fig 7). Few loci had antagonistic effects alike the QTL V__SBxGN__161933 that influences fermentation kinetics V50_80 according to the grape must. The supplementary file S6 provides a complete overview of GxE for the 78 QTLs detected in this work.

**Fig 7.**
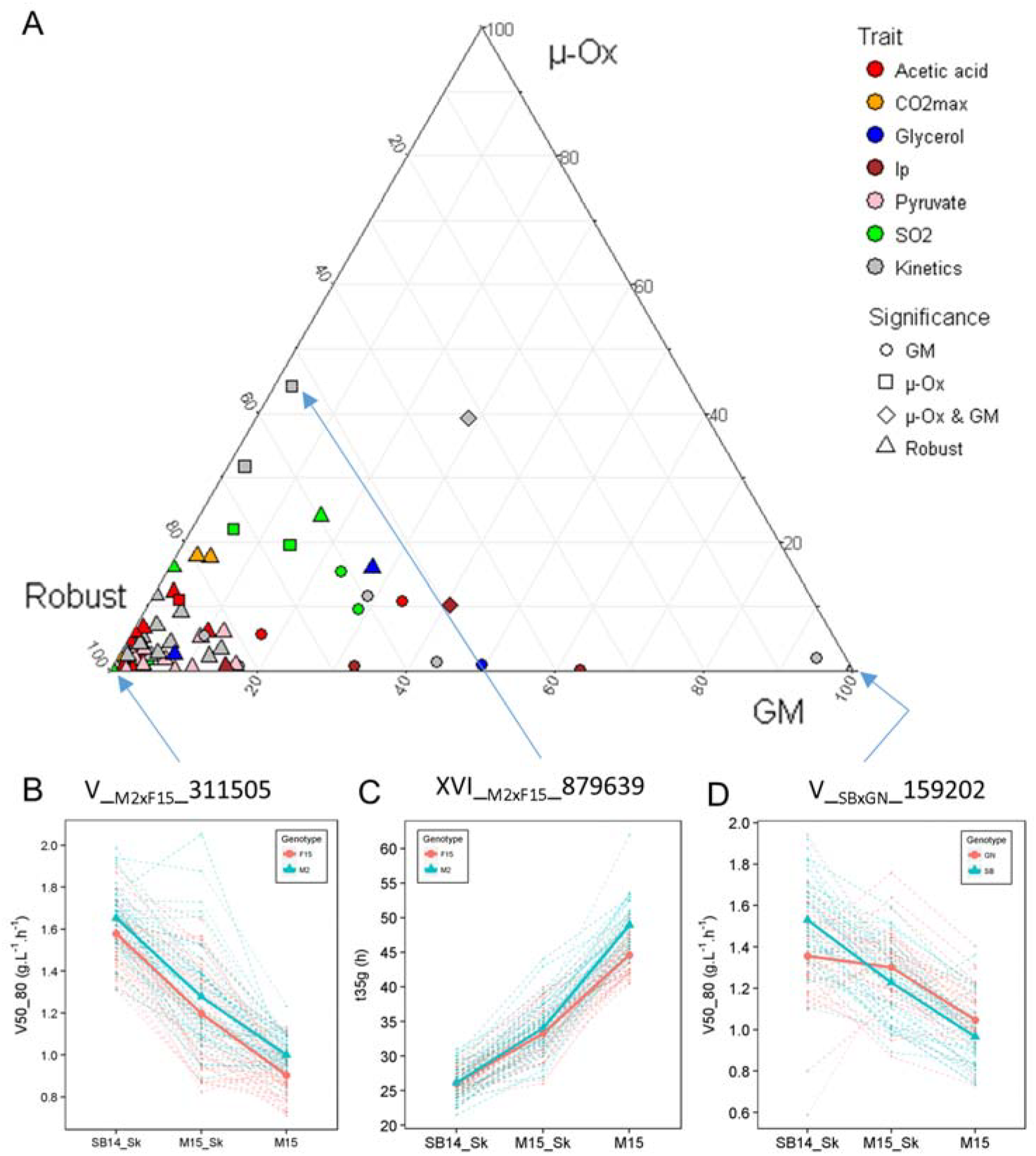
Interaction level of QTLs and environmental conditions. Panel A. The ternary plot shows the proportion between genetic effect (G), interaction with grape must (GM) and interaction with micro-oxygenation (μ-Ox) for each QTL. Significance levels were assessed by ANOVA (α = 0.05). Panel B. Three examples of QTLs for which extreme levels of interaction were identified (i.e. the less interacting QTL, the most interacting QTL with micro-oxygenation and the most interacting QTL with grape must). Dashed lines show the reaction norm of each segregant according to their allele inheritance. Full lines show average value of the all the segregant according to marker inheritance.

### Molecular dissection of GxE interactions

The accuracy of the QTL mapping and the powerful molecular genetics of *S. cerevisiae* offer the possibility to bridge the gap between a QTL and the causative nucleotide variations. Two particular GxE interactions are discussed in this section. Among the interactive QTLs, the locus XV__M2xF15__26284 interacts with both GM and μ-Ox, impacting fermentation kinetics (S5 Fig, panel A). The inheritance of the F15 allele increases *V50_80* only in M15_Sk conditions (S5 Fig, panel B). This *GxE* effect is strong enough to shape a great part of the norm of reaction of M2xF15 offspring. Indeed, k-mean clustering applied to M2xF15 progeny clones defined two non-parallel reaction norm groups. The first one was significantly enriched in strains that inherited the F15 allele and the second one in strains that inherited the M2 allele (CHI2 test, α < 0.05) (S5 Fig, panel C). Although the marker was localized in the gene named *HXT11*, an accurate analysis of the parental sequences revealed that strain F15 doesn’t have this gene. Interestingly, many industrial strains show a deletion of this region (Dunn, Levine, et Sherlock 2005). A copy number variation (CNV) analysis of this region demonstrates that F15 lacks five ORFs (*CSS3, ENB1, IMA2, HXT11*), which are present in the other parent M2 (S5 Fig, panel D). The absence of one of these genes should explain the phenotypic discrepancy observed.

The second marker was XVI__SBxGN__373847, located in the gene *SSU1* that strongly linked three traits (*lp, SO*_*2*_, and *V50_80*). Since this QTL was physically linked to the marker XV__SBXGN__172951 we concluded that the XV-t-VXI translocation could play a pleiotropic role during alcoholic fermentation. In addition, this QTL showed important *GxE* interactions with GM that are likely due to the difference in the SO_2_ effect between red and white grape juices. In order to test if the *SSU1* inheritance has both pleiotropic and GxE effects, we compared the phenotypic response of previously obtained hemizygous hybrids. Those hybrids are isogenic to SBxGN but have only one functional copy of *SSU1*. The allele *SSU1*^*SB*^ is located in a wild type chromosomal environment (XVI-wt), while the allele *SSU1*^*GN*^ has a translocated environment that increases its expression due to the proximity of the *ADH1* promoter (Zimmer et al. 2014). The phenotypic responses of hemizygous hybrids were measured in the three conditions (S7 Table) and compared to the two groups of SBxGN according to their inheritance for the *SSU1* locus (Fig 8). For each of the traits investigated, the progenies reaction norms and hemizygous hybrid responses were very similar. Both interacted significantly with GM and the slope on variation was oriented in the same direction. This result demonstrates that *SSU1* allele inheritance accounted for the main part of the non-parallel reaction norm of the SBxGN progeny. Moreover it demonstrates the pleiotropic effect of the *SSU1* gene that impacts lag phase duration but also unrelated phenotypes such as end-product concentration of SO_2_ as well as the fermentation kinetics.

**Fig 8.**
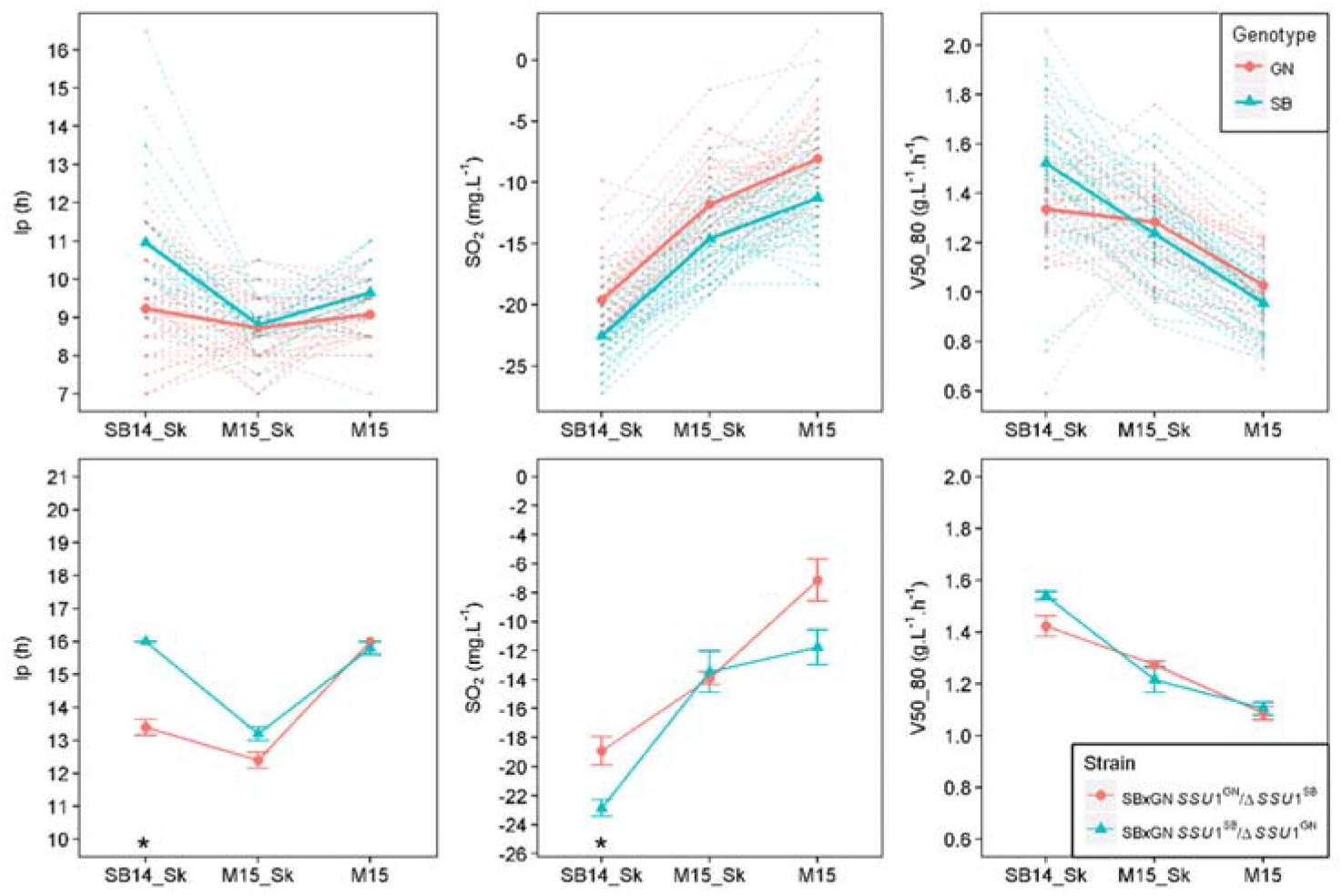
Effect of the XV-t-VXI translocation on phenotypic plasticity. Panel A. The norm of reaction of the segregants that inherited from SB or GN for QTL XVI__SBxGN__373847. Dashed lines show the reaction norm of each segregant according to their allele inheritance. Full lines show average value for all segregants according to marker inheritance. Panel B. The norm of reaction of the hemizygotes. A star means significant difference (Wilcoxon, α = 0.05).

## Discussion

This work aimed to estimate and identify the genetic determinism of phenotypic plasticity. In the last decades, this universal phenomenon has moved from a marginal interest to a new paradigm shaping the evolution of living species in their biotopes (West-Eberhard 1989; Pigluicci 2005; Nussey D. H., Wilson a. J., et Brommer J. E. 2007). Although of great interest, molecular examples of natural genetic variations having a relevant GxE effect are quite scarce (Gerke et al. 2010; Campitelli, Des Marais, et Juenger 2016; Martí-Raga et al. 2017). To challenge this task, we used the species *S. cerevisiae* as it is a powerful tool for achieving quantitative genetics (Liti et Louis 2012). Winemaking conditions allowed us to reproduce a complex and changing environment suitable for identifying alleles conferring phenotypic plasticity. Therefore, in order to match enological practices as closely as possible, the genetic material used was derived from commercial wine starters and the culture conditions used were natural grape juices. This choice was also motivated by the possibility of using the QTLs detected in further selection programs using molecular markers as previously described (Marullo et al. 2007; Dufour et al. 2013; Noble, Sanchez, et Blondin 2015).

### Lessons from QTL mapping with divergent wine parental pairs

Before achieving the QTL analysis, we compared the two population progenies demonstrating that the parental pairs are quite divergent. Indeed, in the SBxGN cross the genetic and phenotypic distance of parental strains was much more important than in the M2xF15 one. In yeast breeding, the selection of parents is mostly based on their phenotypic values. Indeed for optimizing numerous traits, the common strategy consists of selecting parental strains having extreme and opposite phenotypes in order to combine their allelic set in a unique strain (Marullo et al. 2006). This rational has been adopted by many authors for achieving QTL mapping programs (Ambroset et al. 2011; Brice et al. 2014; Jara et al. 2014; Martí-Raga et al. 2017). For the first time, this work compares the efficiency of QTL mapping performed with hybrids resulting from akin and distant yeast strains. Surprisingly, QTL mapping efficiency was similar in both populations since the number of QTLs detected as well the part of variance they explained were very similar (Fig 6 and S6 Fig). Another noteworthy result is the near absence of QTL co-localization (only two loci for 78 QTL detected). This finding has been previously reported in *S. cerevisiae* where the majority of the QTLs are specific to a single cross-combination (Cubillos et al. 2011). However, the four parental strains used in that study were widely divergent (African or Malaysian forest, sake fermentation (Japan) and European wine). In the present work, we observed similar results with strains derived from the same biotope (wine fermentation) that have been subjected to intensive human selection (commercial starters). This means that most yeast strains underwent numerous mutations in different loci affecting the same phenotype, as previously proposed (Liti et Louis 2012). This observation was supported by the fact that phenotypic segregation in both offspring generated phenotypic variability exceeding those found in a wide panel of commercial starters. This offers the possibility of improving strain performance by tapping into the natural mutation pool present within the population of wine strains. However, each have a reduced effect, less than 10% of the explained variance, which is consistent with the infinitesimal model (Lynch et Walsh 1998, 1:198). The use of two divergent populations was then beneficial for capturing more genetic variability, multiplying by two the potential number of natural variation to explore. This also underlines the wide number of mutations in the yeast genome that can affect a phenotype. The expressivity (penetrance) of such mutations in different backgrounds should be very low due to underlining epistasis. However, two common QTLs were found suggesting that they could be due to positive ongoing selection as discussed below.

### Phenotypic plasticity is the rule and QTL mapping allows the capture of its genetic determinism

In order to detect QTLs explaining plasticity, eleven quantitative traits were measured in three environments. The conditions applied were chosen for reflecting two relevant enological parameters, the nature of grape must (GM) and the micro-oxygenation (μ-Ox) (Peltier et al. 2018). We first characterized the reaction norm of all the strains, demonstrating that an important part of observed variance was due to strain x environment interactions (up to 26 %, Table 1). The acetic acid patterns measured for 189 strains (Fig 5) revealed that non-parallel reaction norms are the rule and not the exception. By changing the oxygen input (from 2 mg.L^−1^ in the unshaken condition to 4 mg.L^−1^ in the shaken condition) or by changing the grape must, a variability was observed for 96.3 % of the strains with amplitudes that are relevant for the wine industry. As for QTL detection efficiency, the hybrid genetic background does not seem to govern those patterns. This original result demonstrates that GxE are important in wine fermentation and should be better investigated.

The multi-environment QTL analysis drastically increased the statistical power of QTL detection since most of the QTL (45) were only identified with this model, confirming the efficiency of this strategy as previously described (Jiang et Zeng 1995; Chen, Zhao, et Xu 2010; El-Soda et al. 2014). Five QTLs were only identified with a single-environment model. Rather than original QTLs, these peaks seem to be QTLs with ambiguous positions that vary greatly between single and multi-environment analyses. Indeed, in an extended window of 100 kb, a QTL identified with the multi-environmental model was found for the same trait. The use of several environments makes it possible to evaluate QTLs robustness. Here, 72 % of the QTLs are statistically robust against environment. Since the applied conditions are very different from an enological point of view (white vs red grape must, hypoxic vs micro-oxygenation), most of these QTLs should be robust for most of the fermentations and are suitable for developing selection programs assisted by molecular markers. The 28 % of remaining QTLs had a significant interaction with the environment. Except for two QTLs showing antagonistic effects, the interactions detected were due to “scale effect”. This contrasts with the observation made by (A. Bhatia et al. 2014) where QTLs of interaction mostly had antagonistic effect according to environment or were specific to one environment. However the various media tested in that work were quite divergent since the yeast growth was measured in either fermentable or non-fermentable carbon sources, thus creating drastic physiological switches. In our study, the conditions applied remain restricted to the fermentation creating fine-grained interactions.

This study introduces an approach to bridge the gap between non-parallel reaction norms observed and their underlying genetic causes. Some QTLs with significant GxE effects were matched with the strain reaction norms. Among the nine QTLs with significant GxE effects for kinetic traits, two of them (XV__M2xF15__26284 and XVI__SBxGN__371802) had high interaction levels explaining 3.7 % and 5 % of the variance, respectively. Their inheritance shaped a large part of the phenotypic plasticity of the *V50_80* in both populations (Fig 8, panel C and S5 Fig, panel C) separating the strains in the distinct reaction norms predicted by the k mean clustering (S2 Fig). When non-parallel reaction norms were more complex alike *acetic acid* and *pyruvate*, the few QTLs with significant GxE interactions were not sufficient for explaining the overall phenotypic response observed (Fig 5). However those loci could be helpful for developing breeding programs focusing on specific applications. For example, some strains do not have the appropriate phenotypic response in the presence of micro-oxygenation by not decreasing their acetic acid production (Fig 5). By interacting with μ-Ox, the QTLs V__SBxGN__70702 and XIV__SBxGN__623501 explained a part of this phenotypic response. The incorporation of their favorable alleles by a breeding strategy in other backgrounds, which show inappropriate response to micro-oxygenation, could be very valuable. Their additive effect would reduce 66 % of these compounds in micro-oxygenated conditions. However, all the phenotypic responses were not explained. Indeed, none of the QTL identified for *acetic acid* explains the phenotypic response of strains of cluster GM and μ-Ox that had an increased production of acetic acid in SB14_Sk compared to M15_Sk (Fig 5). Similarly for *pyruvate*, no interactive QTL was identified while highly divergent norm of reaction were obtained for this trait. This large number of type of reaction norms may reflect a complex genetic determinism with a high number of interacting loci. The segregation of these factors in less than a hundred individuals does not allow their identification. Therefore, the characterization of the genetic determinism of complex traits such as that of *acetic acid* or *pyruvate* seems to require the study of a larger population

### The promoter region of the gene SSU1, a genetic locus under balanced selection that have pleiotropic effects

Within an evolving population subject to selection, the genetic variations are a mix of three types: (i) rare and deleterious alleles resulting from recent mutational events not yet eliminated by selection; (ii) neutral alleles whose frequencies follow the rules of genetic drift; (iii) alleles with intermediate frequencies that are subjected to different counterbalanced selection pressures (Robertson 1956; Turelli et Barton 2004; Mackay 2010). For this last type of allelic variation, the selection pressure can be balanced because one allele may have a favorable or an unfavorable effect according to the environment in which it is expressed, meeting the QTLs described by (Aatish Bhatia et al. 2014). Otherwise, the selection pressure can be balanced because one allele can have a pleiotropic effect on fitness parameters, positively affecting some traits but negatively other ones creating phenotypic trade-offs. Mutations with balanced effects have already been identified in *Arabidopsis thaliana* for water use efficiency (Campitelli, Des Marais, et Juenger 2016) or delay of germination (Chiang et al. 2011). They have been useful for understanding why particular accessions are better adapted to specific environment. In yeast, pleiotropic genes (*MKT1 or IRA2*) that affect unrelated traits like temperature resistance (Steinmetz et al. 2002; Parts et al. 2011) or sporulation efficiency (Ben-Ari et al. 2006) were previously described. The effect of the gene *SSU1* in the context of the translocation XV-t-XVI is an interesting case where a single gene can have pleotropic effects on several phenotypes and interaction with environments.

In *S. cerevisiae*, the sulfite pump *Ssu1p* is required for efficient sulfite efflux (Park et Bakalinsky 2000). The expression of this transporter can be strongly enhanced by chromosomal translocations that modify the promoter environment of *SSU1*. This enhanced expression confers a higher resistance to the inhibitory effect of SO_2_ added to the grape must. Two independent translocations (VIII-t-XVI and XV-t-XVI) have been reported in the literature (Pérez-Ortín et al. 2002; Zimmer et al. 2014), and are hallmarks of adaptation to winemaking practices (Marsit et al. 2017). Both translocations confer an important adaptive advantage in respect to indigenous flora by reducing the lag phase duration, however, they are not present in all wine strains and are often present in a heterozygous state (Nardi et al. 2010; Zimmer et al. 2014). In this work, we demonstrate that both translocations impact the final amount of SO_2_ and the late fermentation rate. Using hemizygous hybrids previously constructed, we validated the pleotropic effect of *SSU1* in the SBxGN background (Fig 8). The phenotype of the translocated form is in agreement with the expected increase in *SSU1* expression. Therefore, translocated progeny clones leave more SO_2_ at the end of fermentation. More startling is the translocation effect on the late fermentation rate (*t80g, V80_80*). Indeed, in the SB14_Sk conditions (containing more active sulfites), the translocated strain has a slower end-fermentation rate due to a relative lower viability (data not shown) due by the higher SO_2_ concentration still present in the medium. This opposed effect of yeast fitness (short lag phase but lower fermentation rate and viability) constitutes a phenotypic trade-off and suggests that *SSU1* has undergone a balanced selection. Despite the great advantage conferred by a short lag phase (more than 20 hours in high sulfite conditions) the non-translocated form of *SSU1* also confers adaptive advantages. Indeed, due to their low SO_2_ efflux, the strains having the non-translocated allele do not have to cope with the toxicity of SO_2_ in the late stages of the fermentation, which improves yeast viability. Moreover the effect of the translocation on lag phase duration is much lower in red grape must (Merlot) creating environmental conditions that preserve this allele from selective pressure. Pleiotropic genes are likely important levers of the complex architecture of quantitative traits and they have been reported in other living organisms. The gene *SSU1*, with its translocated forms, is a good example of pleotropic gene promoting phenotypic trade-offs conditioned by environmental conditions.

## Materials and methods

### Yeast strains and culture media

Strains used in this study belong to the yeast species *Saccharomyces cerevisiae*. The four parental strains (SB, GN, M2, F15) are diploid homothallic monosporic clones derived from wine commercially available starters. The strains SB, GN and F15 were derived from VL1, Actiflore BO213, Zymaflore F15 (Laffort, Bordeaux, France), respectively, while M2 was derived from Oenoferm M2 (Lallemand, Blagnac, France). The two populations used to perform QTL mapping were obtained from two F1-hybrids (M2xF15 and SBxGN). The first offspring (95 individuals) was derived from the F1-hybrid M2xF15 generated by (Huang, Roncoroni, et Gardner 2014). The second progeny (94 individuals) was obtained by tetrad dissection of the F1-hybrid SBxGN (formerly named HO-BN by (Marullo et al. 2007)). Both progeny clones are homozygous and diploid due to the homothallic nature of all the parental strains. All the strains were grown at 28 °C on YPD medium (1 % yeast extract, 1 % peptone, 2 % glucose), solidified with 2 % agar when required. Sporulation was triggered by plating fresh cells on potassium acetate medium after three days at 24 °C. The strains were stored long term in YPD with 50 % of glycerol at −80 °C. Construction of the hemizygotes SBxGN_SSU1^SB^/ΔSSU1^GN^ and SBxGN_SSU1^GN^/ΔSSU1^GB^ (formerly named G092G and S092S) has been described by (Zimmer et al. 2014).

### Fermentations

The two grape juices used, Merlot of vintage 2015 (M15) and Sauvignon Blanc of vintage 2014 (SB14), were provided by *Vignobles Ducourt* (Ladaux, France) and stored at −20°C. Before fermentation, grape juices were sterilized by membrane filtration (cellulose acetate 0.45 μm Sartorius Stedim Biotech, Aubagne, France). Their main enological characteristics are given in Table 2. Sugar content, assimilable nitrogen, pH, total and free SO_2_ were assayed by the enological analysis laboratory (SARCO, Floirac, France). Malic acid was determined by enzymatic assay (Peltier et al. 2018).

**Table 2.** Grape juices used in the study.

| Grape must | Code | Sugar content (g.L<sup>-1</sup>) | Assimilable Nitrogen (mg N.L<sup>-1</sup>) | Malic acid (g.L<sup>-1</sup>) | pH | Total SO<sub>2</sub> (mg.L<sup>-1</sup>) | Free SO<sub>2</sub> (mg.L<sup>-1</sup>) | Molecular SO<sub>2</sub> (mg.L<sup>-1</sup>) |
| --- | --- | --- | --- | --- | --- | --- | --- | --- |
| Sauvignon Blanc 2014 | SB14 | 194 | 157 | 5.6 | 3.19 | 34 | 7 | 0.32 |
| Merlot 2015 | M15 | 219 | 99 | 1.9 | 3.53 | 46 | 33 | 0.68 |

Fermentations were carried out as previously described by (Peltier et al. 2018). Briefly, fermentations were run at 24°C in 10 mL screw vials (Fisher Scientific, Hampton, New Hampshire, USA) with 5 mL of grape must. Hypodermic needles (G 26 – 0.45 × 13 mm, Terumo, Shibuya, Tokyo, Japan) were inserted through the septum for CO_2_ release. Two micro-oxygenation conditions were used by applying or not constant orbital shaking at 175 rpm during the overall fermentation. During this study, three fermentation conditions were used: *SB14* with shaking (*SB14_Sk*), *M15* with shaking (*M15_Sk*) and *M15* without shaking (*M15*).

Fermentation progress was estimated by regularly monitoring regularly the weight loss caused by CO_2_ release using a precision balance. The amount of CO_2_ released over time was modeled by local polynomial regression fitting with the R-loess function setting the span parameter to 0.45. Seven parameters were extracted from the model:

*Ip*(h): the lag phase time observed before to release of CO_2_ at 2 g.L^−1^;

*t35g, t50g* and *t80g* (h): time (minus *lp*) until 35, 50 and 80 g.L^−1^ of CO_2_ were released;

*V15_50* (g.L^−1^.h^−1^): average sugar consumption between 15 % and 50 % of *tCO2max*;

*V50_80* (g.L^−1^.h^−1^): average sugar consumption between 50 % and 80 % of *tCO2max*;

*CO2max*: maximal amount of CO_2_ released (g.L^−1^).

### Metabolic compounds

At the end of the fermentation the concentration of four compounds was measured at the metabolomics platform of Bordeaux (http://metabolome.cgfb.u-bordeaux.fr) by semi-automated enzymatic assays (Peltier et al. 2018). Four phenotypes were measured: acetic acid (g.L^−1^), *glycerol* (g.L^−1^), *pyruvate* (mg.L^−1^) (from the final samples taken from each fermentation) and *SO*_*2*_ *Yield* (mg.L^−1^) ([*SO*_2_]_*final*_ - [*SO*_*2*_]_*initial*_).

Each fermentation was carried out two times for the progeny clones and their respective hybrids and 10 times for each parental strain (M2, F15, SB, GN). The entire data set is given in the S2 Table.

### Genotyping and marker map construction by high throughput sequencing

The procedure used for genotyping the 94 SBxGN progenies was the same as published by (Martí-Raga et al. 2017). Briefly, all the 94 diploids progeny clones were genotyped by whole-genome sequencing at a low coverage (3–6 X). DNA libraries were pooled and sequenced with a MiSeq apparatus using the standard kit v2 (Illumina) generating paired-end reads of 2 × 250 bp. Filtering and mapping of all sequencing data was performed using publicly available tools (https://usegalaxy.org).. Sequencing data were treated as single reads. The main parameters of filtering and mapping were: read trimming (−39 bases), Phred quality cut-off (Q=20), and read mapping (BWA software with default parameters). Once the reads had been mapped, BAM files were extracted and a *pileup* dataset was generated using SAMTools’ (Li et al. 2009) for every segregant. The *pileup* dataset was opened in R and SNP between the parental strains was evaluated using an R script (Martí-Raga et al. 2017). To construct the marker map, we retained the markers with a 1:1 segregation among the progeny (Chi-x test, α > 0.05) and being evenly distributed along the genome (1 marker/15 kb). The final map generated had 3,433 markers (S2 Table). The procedure used for genotyping the 95 M2xF15 progenies was described by (Roncoroni 2014).

### Microsatellite genotyping

The DNA of *S. cerevisiae* strains was quickly extracted in 96-well microplate format using a customized LiAc-SDS protocol (Albertin et al. 2017). Fifteen polymorphic microsatellite loci SCAAT3 (*C3, C5, SCYOR267C, C8, C11, SCAAT2, YKL172, SCAAT6, C9, C4, SCAAT5, SCAAT1, C6, YPL009, YKL172W*) were used for estimating the genetic relationships within 96 commercial starters and the four monosporic parental strains (GN, SB, M2, F15) used in this work (S3 Table). The genotyping conditions used were broadly those described by (Börlin et al. 2016). Briefly, two multiplex PCRs allowing genotyping of seven loci were carried out in a final volume of 12.5 μL containing 6.25 μL of the Qiagen Multiplex PCR master mix and 1 μL of DNA template, and 1.94 μL of each mix was added in the mixture using the concentrations indicated in the S9 Table. Both reactions were run with the following program: initial denaturation at 95 °C for 5 min, followed by 35 cycles of 95 °C for 30 s, 57 °C for 2 min, 72 °C for 1 min, and a final extension at 60 °C for 30 min. The size of PCR products was analyzed by the MWG company (Ebersberg, Germany) using 0.2 μL of 600 LIZ (GeneScan) as a standard marker. Chromatograms were analyzed with the GeneMarker (V2.4.0, Demo) program.

### Data analyses

All the statistical and graphical analyses were carried out using R software (R Development Core Team 2011).

### Estimation of environment, cross and strain effect

The variation of each trait was estimated by the analysis of variance (ANOVA) using the *aovp* function of the *lmPerm* package in which significance of the results were evaluated by permutation tests instead of normal theory tests.

The *LM1* model estimated the effects of the cross, of the environment and of the cross-by-environment interaction of fermentation traits according to the following formula:

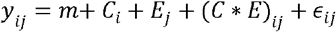

where *y*_*ij*_ was the value of the trait for cross *i* (*i* = 1, 2) in environment *j* (*j* = 1, 2, 3), *m* was the overall mean, *C*_*i*_ was the cross effect, *E*_*j*_ the environment effect, (*C* * *E*)_*ij*_ was the interaction effect between cross and environment and *ε*_*ijk*_ the residual error.

The *LM2* model estimated the effects of the strain, of the environment and of the strain-by-environment interaction on fermentation traits according to the following formula:

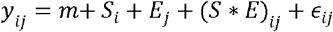

where *y*_*ij*_ was the value of the trait for strain *i* (*i* = 1, …, 189) in environment *j* (*j* = 1, 2, 3), *m* was the overall mean, *S*_*i*_ was the strain effect, *E*_*j*_ the environment effect, (*S * E*)_*ij*_ was the interaction effect between strain and environment and *ε*_*ijk*_ the residual error.

### Estimation of the genetic distances within strains

The microsatellite dataset was manipulated using the *adegenet* package (Jombart 2008) implemented in R. The percentage of missing data was 1.6 %. The genetic distance within the strains was estimating using the Bruvo’s distance using the *poppr* package (Kamvar, Tabima, et Grünwald 2014). The unrooted dendrogram was built by Neighbor Joining (*ape* package) (Paradis, Claude, et Strimmer 2004). Since the bootstraps estimated did not allow the resolution of clear groups, the genetic structure was estimated by a *k* mean clustering using the function *find.cluster* allowing the detection of three main groups. The minimal goodness of fit was selected using comparing AIC, BIC and WSS criteria using default parameters.

### Norm of reaction clustering

For each trait the distance between norms of reaction were calculated according to the formula:

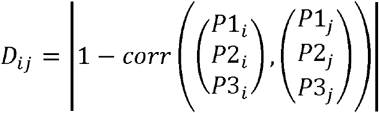

where *D*_*ij*_ was the distance between strains *i* and *j*, *P*1_*i*_ and *P*1_*j*_ the phenotypic values measured in *SB14_Sk*, *P*2_*i*_ and *P*2_*j*_ the phenotypic values measured in *M15_Sk* and *P*3_*i*_ and *P*3_*j*_ the phenotypic values measured in *M15* for strains *i* and *j*, respectively. The minimal distance (0) was set between norm of reactions with null variances and the maximal distance (1) was set between norm of reaction with a null variance and all the other norm of reactions with a non-null variance. K-means clustering was performed on distance matrix with the *pam* function of the *cluster* package of the R program. Appropriate number of cluster was determined by the best silhouette value between one and ten clusters with the *silhouette* function of the *cluster* package of the R program.

### Estimation of heritability, transgression level and phenotypic distance within segregating population

The *lato sensu* heritability *h*^*2*^ was estimated for each phenotype according to (Marullo et al. 2006) as follows:

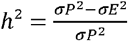

where *σP*^2^ is the variance of progeny population in each environmental conditions, explaining both the genetic and environmental variance of the phenotype measured, whereas *σE*^2^ is the median of the variance of replicates in each environmental conditions, explaining only the environmental fraction of phenotypic variance.

Percentage of transgression was calculated as described by (Marullo et al. 2006). Results were displayed on a heatmap (*heatmap.2* function) of the *gplots* package (Warnes, G 2009).

### QTL mapping

Before linkage analysis phenotypes were normalized by Rank-transformation using the *GenABEL* package (Aulchenko et al. 2007). The QTL mapping analysis was performed with the *R/qtl* package (Broman et al. 2003) on the data collected in the three environmental conditions by using the Haley-Knott regression model and adding the effect of the environment as additive and interactive covariates. For each phenotype, a permutation test of 1000 permutations tested the significance of the LOD score obtained, and a 5% FDR threshold was fixed for determining the presence of QTLs. QTLs identified for the fermentation rate parameters *t35g t50g t80g V15_50* and *V50_80* being in the same 10 kb windows were considered as a unique locus. The QTL position was estimated as the marker position with the highest LOD score among all markers above the threshold in a 30 kb window.

### Principal component analysis

The phenotypic variability of M2xF15 and SBxGN progenies measured in M15 was visualized by a Principal Component Analysis (PCA) using the *ade4* package. An additional dataset, recently obtained by (Peltier et al. 2018), was added to the projection corresponding to the phenotypic values of 31 commercial strains. M15 condition was compared with the only similar condition described in that work (noSk.5_SV). The contribution of each phenotypes to the PCA dimensions and the correlation circle were obtained by the *fviz_contrib* and the *fviz_pca_var* functions of the *factoextra* package (Kassambara A et Mundt F 2015). Phenotypic distances were computed by calculating the Euclidian distances within the strains of the same group in each environmental condition and for each trait. For comparison of the overall phenotypic distance between the commercial starters (n=31) and each progeny population (n = ~100), 1,000 random pools of 31 spore clones of each progeny population were compared to the commercial dataset by a one way analysis of variance *aovp* function of *ImPerm* package. Tukey’s honest significant difference *post hoc* test was used to confirm differences between groups (α = 0.05).

### Estimation of the level of QTL interaction

The interaction level of each QTL was estimated by ANOVA using the *aovp* function of the *ImPerm* package. The LM3 model estimated the effects of the cross, of the environment and of the QTL-by-environment interaction on traits according to the following formula:

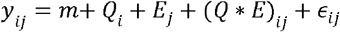

where *y*_*ij*_ was the value of the trait for allele *i* (*i* = 1, 2) in environment *j* (*j* = 1, 2 or *j* = 1, 2, 3), (three combinations of environments were considered: a) SB14_Sk and M15_SK to assess the grape must effect (GM), b) M15_Sk and M15 to assess the micro-oxygenation effect (μ-Ox), c) SB14_Sk, M15_SK and M15 to assess the overall genetic effect), *m* was the overall mean, *Q*_*i*_ was the QTL effect, *E*_*j*_ the environment effect, (*Q * E*)_*ij*_ was the interaction effect between QTL and environment and *ε*_*ijk*_ the residual error. In order to compare the level of interaction across traits and environment, their % of variation explained was calculated by omitting the total sum square of environment. The ratio between grape must interaction, micro-oxygenation interaction and genetic effect was calculated as the ratio of the percentage of variation explained by QxE by considering SB14_Sk and M15_SK for grape must interaction, M15_Sk and M15 for micro-oxygenation interaction and the percentage of variation explained by Q in all conditions for the overall genetic effect.

## Acknowledgments

The authors thank justine Pape, Dylan Dos Reis and Elodie Kaminski that helped managing fermentations. We also thank Warren Albertin for kindly giving the R script used for microsatellite analyses. This work was supported by Région d’Aquitaine.

## Supporting information

**S1 Fig. Meiosis emphases phenotypic novelty**

Panel A. PCA of winemaking properties of M2xF15 and SBxGN progenies and 31 CWS in M15. Panel B. Correlation circle indicating the correlation of the variables for axes 1 and 2. Panel C. Average phenotypic distance computed from M15 condition for CWS, and the two M2xF15 and SBxGN offspring.

**S2 Fig. Clustering of Norm of reaction for each trait**

Norm of reaction of each individual is shown in dotted line and are colored and faceted according their cluster. Solid line shown the average norm of reaction of each cluster. Number of strains within each cluster is indicated by n.

**S3 Fig. Heritability is correlated to the number of QTLs detected**

The data represented are the number of QTLs identified according by cross and by trait according to the average heritability by cross and by trait among the three conditions.

**S4 Fig. Variation of QTL effect according to condition**

For each QTL, the values shown are the difference between the phenotypic values measured for all the segregant that inherited the allele of SB or M2 minus those that inherited from GN or F15. A star means a significant difference (Wilcoxon, α = 0.05).

**S5 Fig. Impact of QTL XV__M2xF15__26284 on V50_80**

Panel A. The norm of reaction of the segregants that inherited from M2 or F15 for QTL XV__M2xF15__26284. Dashed lines show the reaction norm of each segregant according to their allele inheritance. Full lines show average value of the all the segregant according to marker inheritance. Panel B. Enrichment of strains having inherited the F15 allele in cluster 2. Norm of reaction of each individual is shown in dotted line and faceted according to their cluster. M2xF15 progeny clones are colored according to their allele inheritance at XV__M2xF15__26284 (red for F15 and blue for M2). A grey solid line shown the average norm of reaction of each cluster. Number of M2xF15 progeny clones within each cluster is indicated by n=; the first number indicate the number of strains from that inherit from M2 and the second one from strain that inherit from F15. A star means a disequilibrium in the theoretical homogeneous distribution 0.5 / 0.5 chis square test, α = 0.05). Panel C. Coverage difference between M2 and F15 strains within genomic region of QTL XV__M2xF15__26284.

**S6 Fig. Variance explained by QTL according to cross**

Each dot represent a QTL. Bigger points indicate average. There is no significant difference between the two cross (Wilcoxon test pval >0,05).

**S1 Table. 97 commercial strains used**

**S2 Table. Phenotypic dataset**

**S3 Table. LM1 model for the 11 phenotypes with 189 strains and three conditions of fermentation**

**S4 Table. Phenotypic plasticity at the population level**

The first line indicates M15_Sk average phenotypic values and the two other indicate the effect of each environmental parameter: difference between M15_Sk and SB14_Sk (grape must effect), difference between M15_Sk and M15 (micro-oxygenation effect). Significant differences are indicated by * (Wilcoxon test pval <0,05).

**S5 Table. List of QTL identified**

**S6 Table. Number of QTL and their interaction level**

**S7 Table. Hemizygote dataset**

**S8 Table. Genetic map of SBxGN progeny**

