## Supplementary figures and images for "A study of phenotypic plasticity of *Saccharomyces cerevisiae* in natural grape juices shed light on allelic variation under balanced selection"

### Supplementary Materials

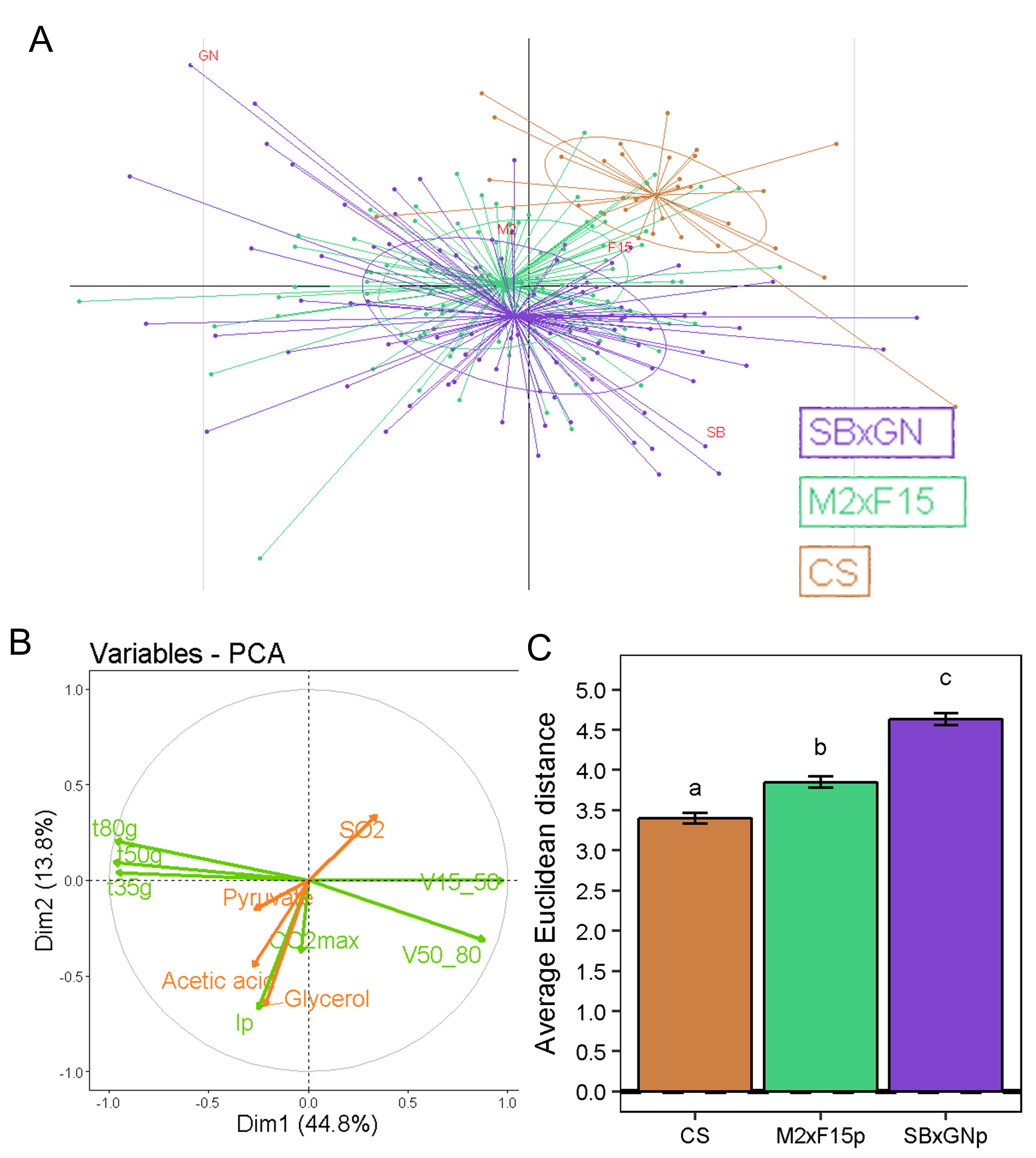

### Supplementary Materials

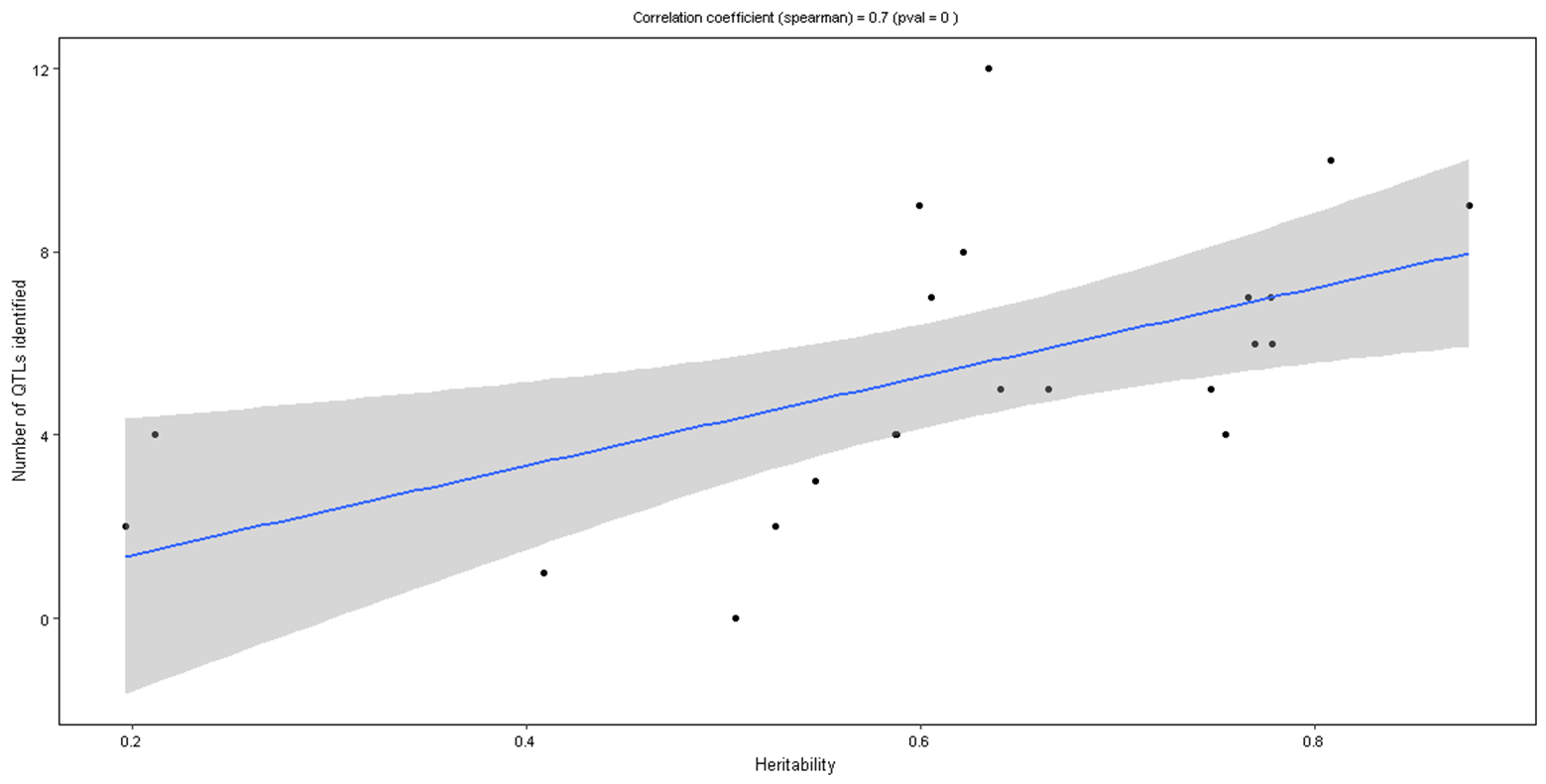

### Supplementary Materials

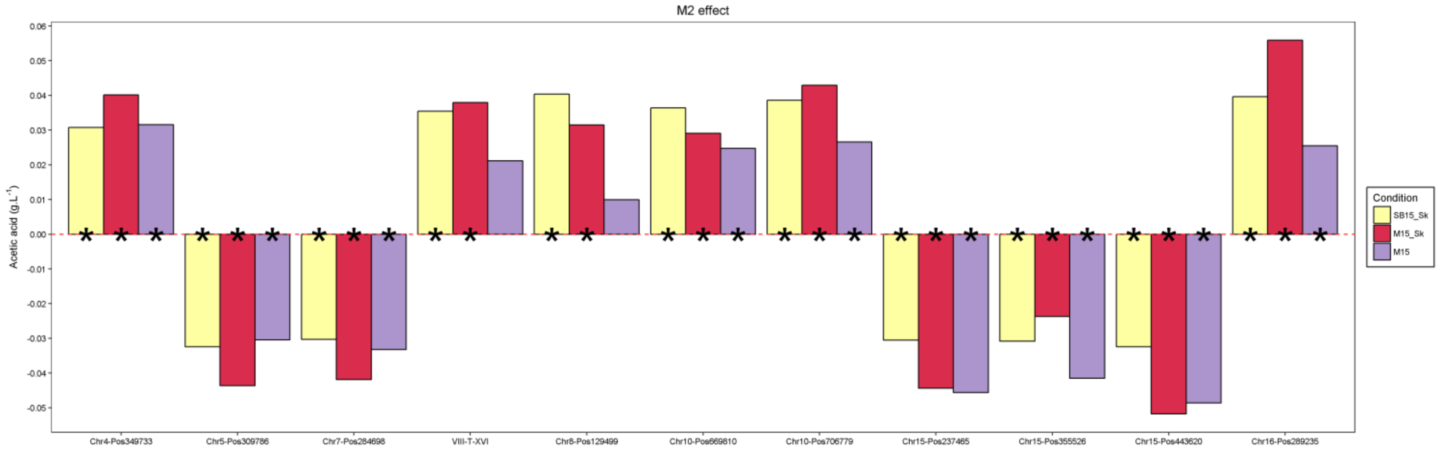

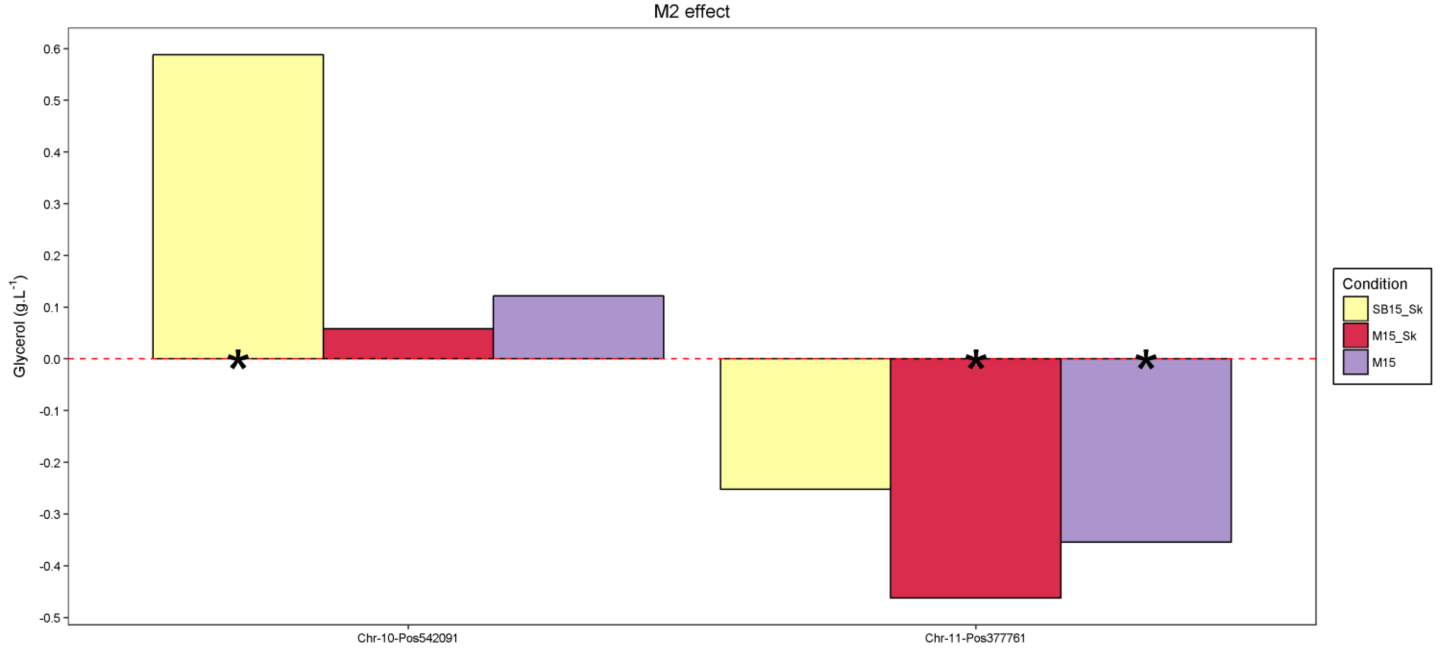

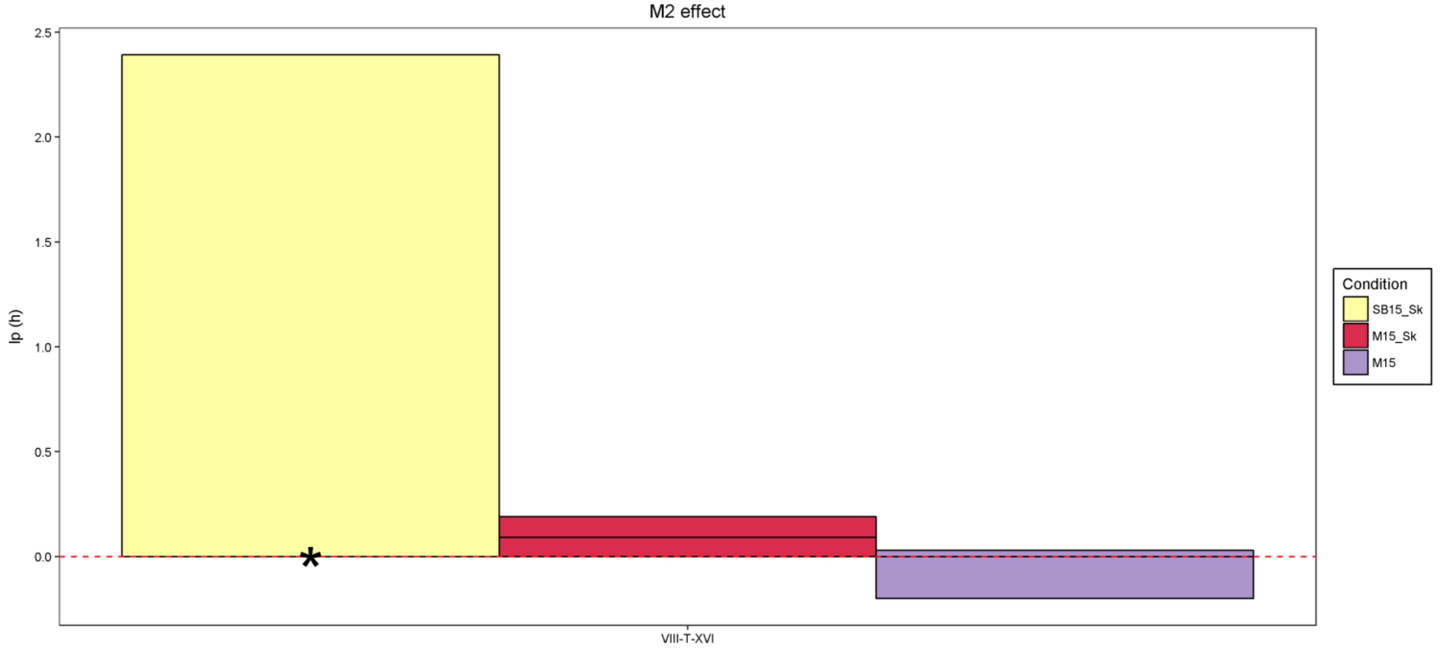

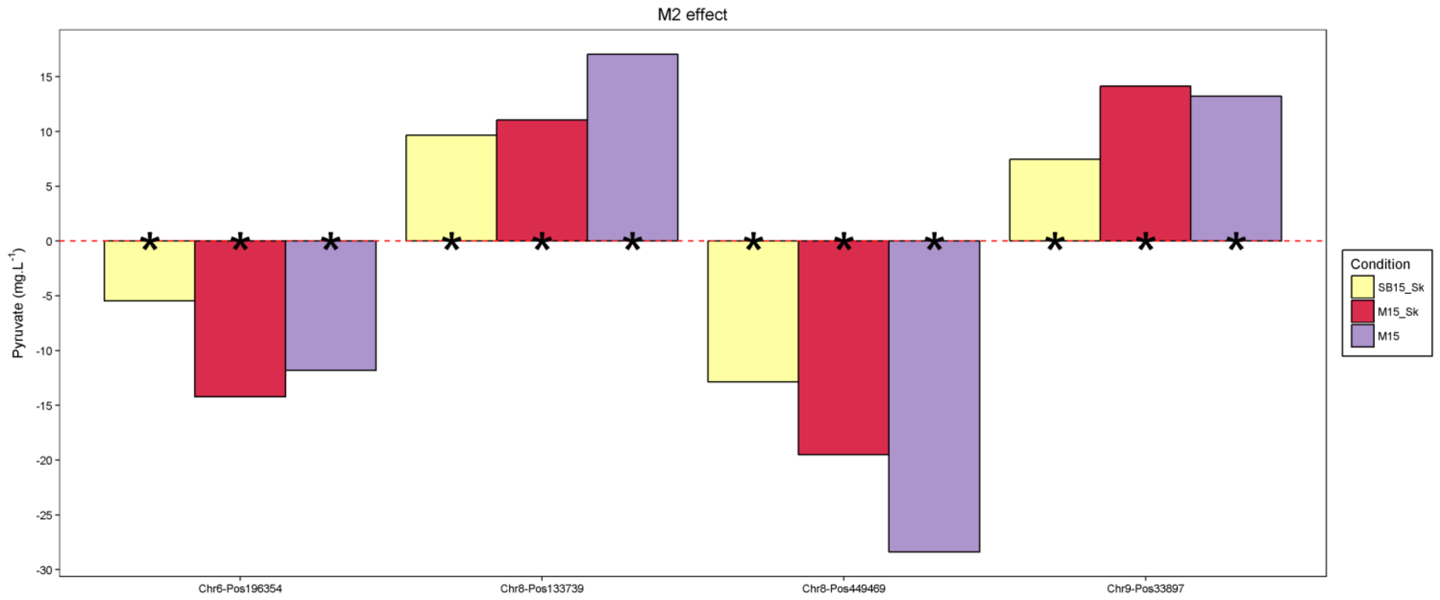

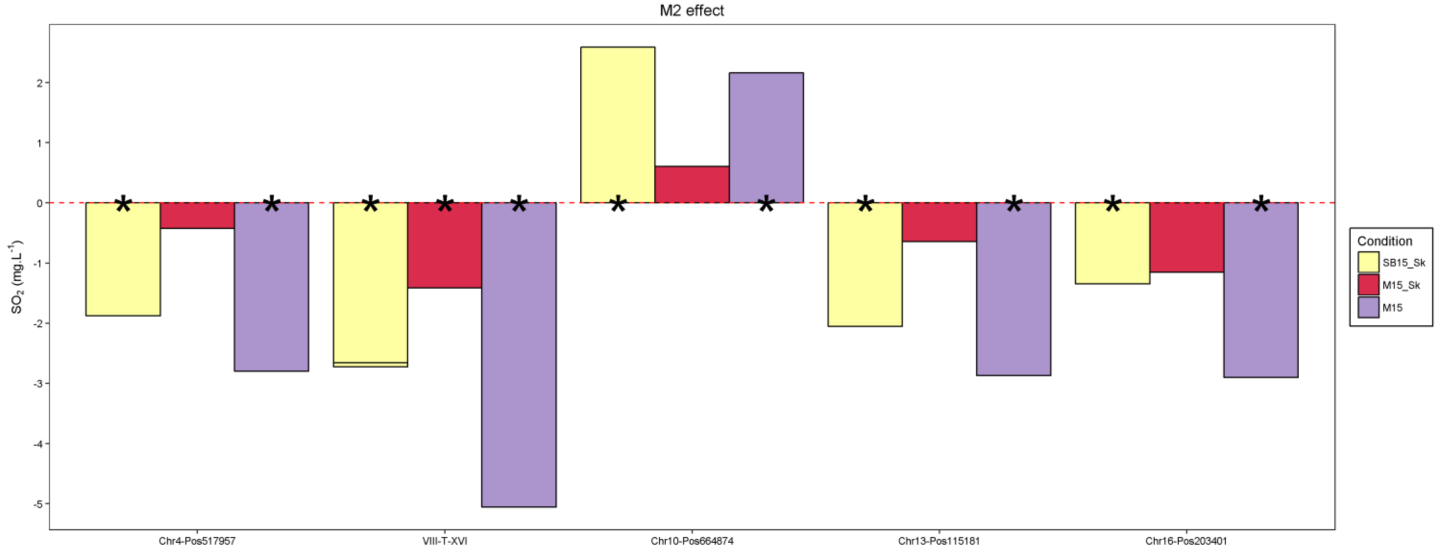

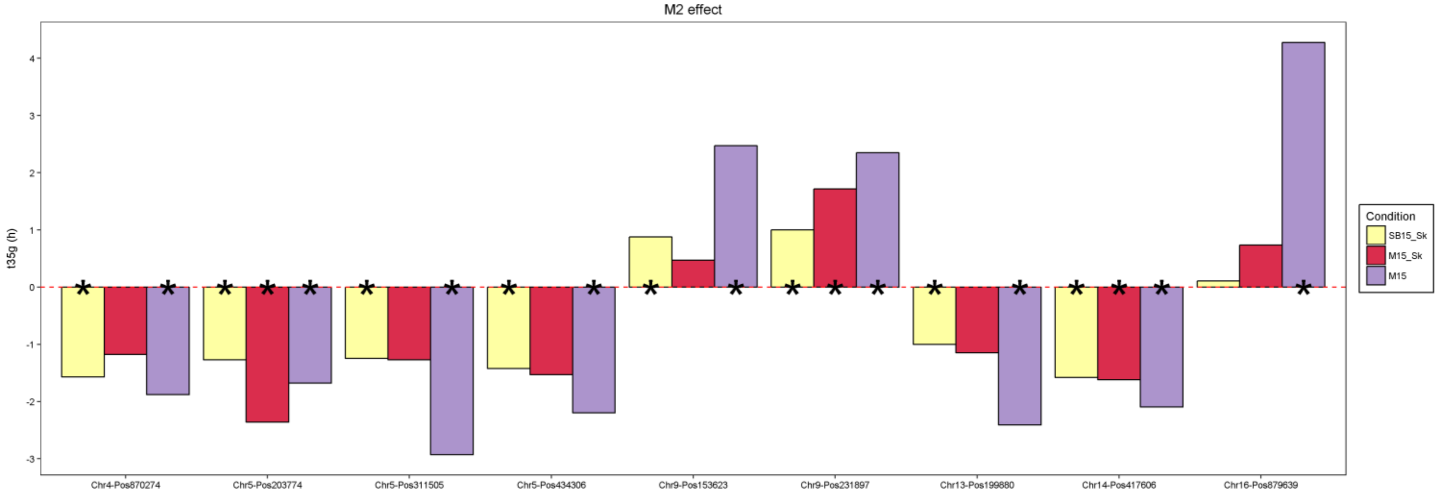

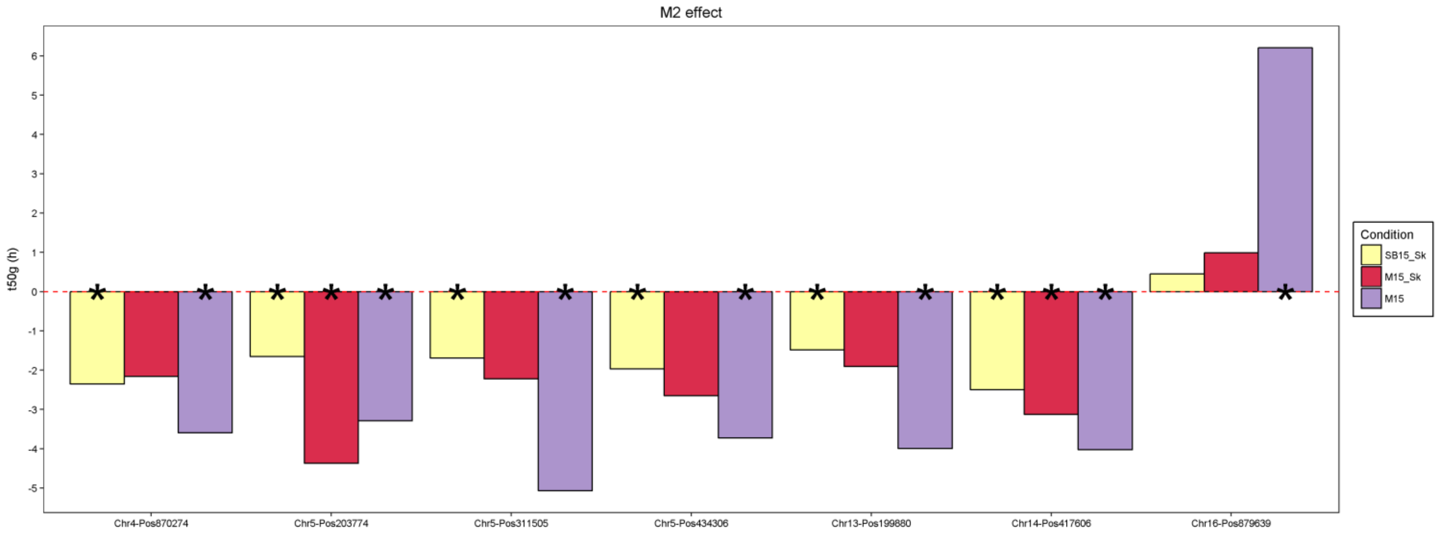

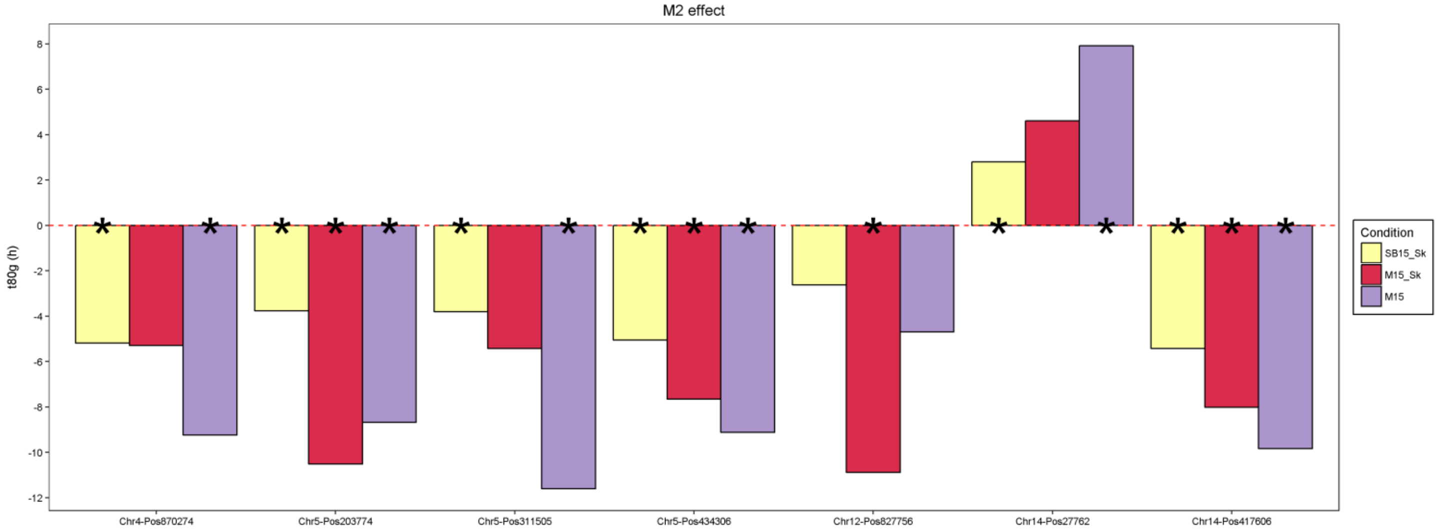

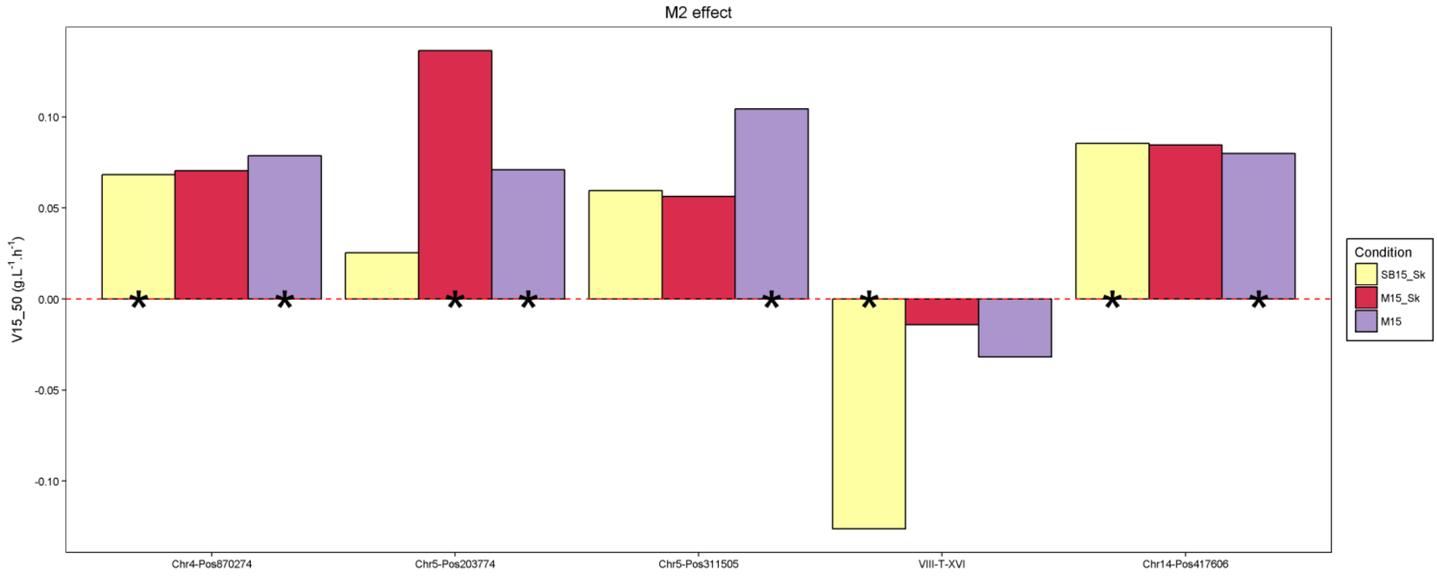

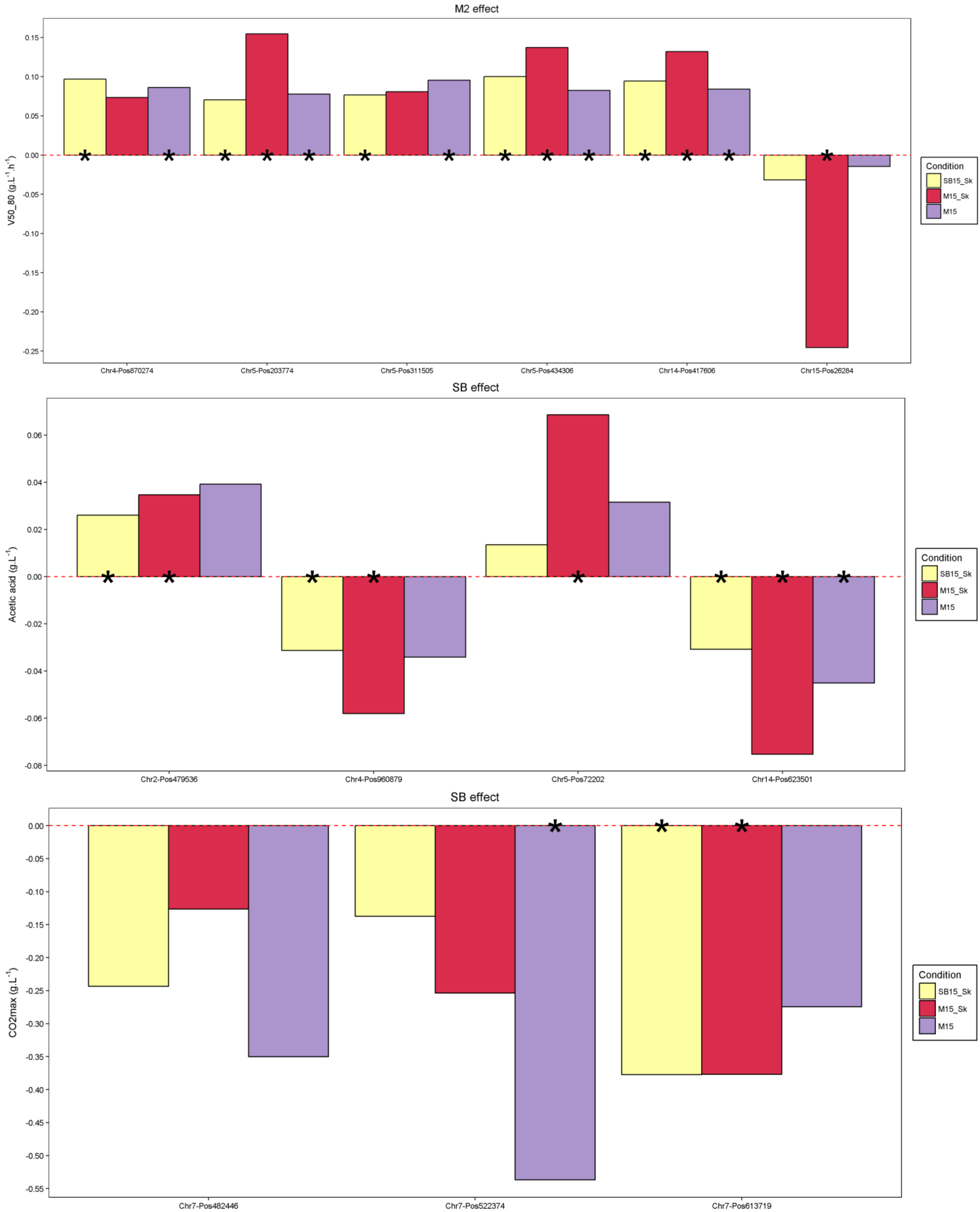

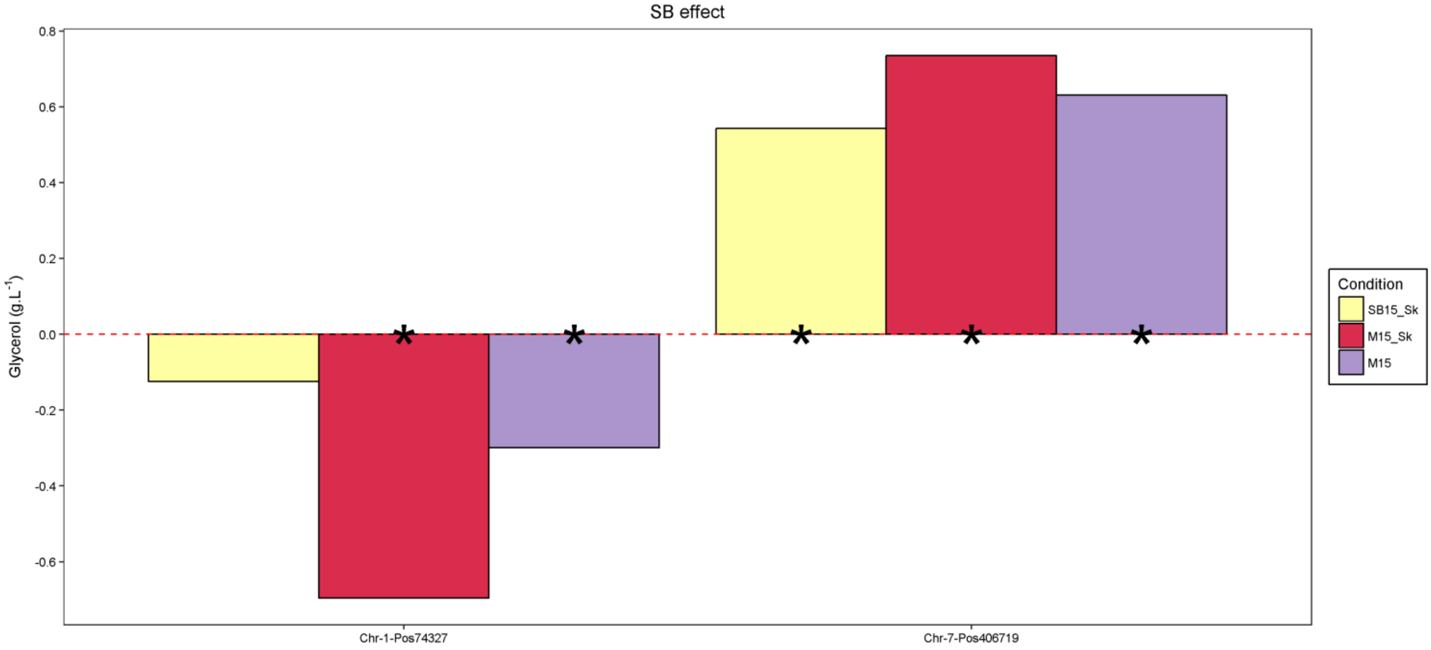

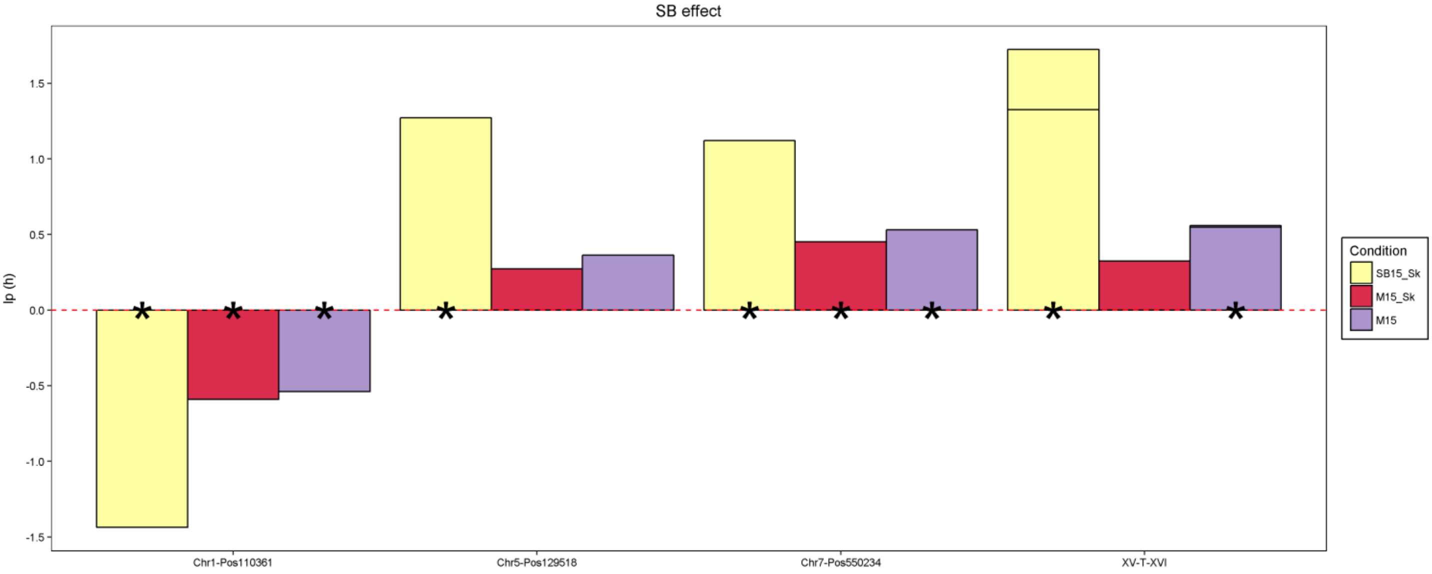

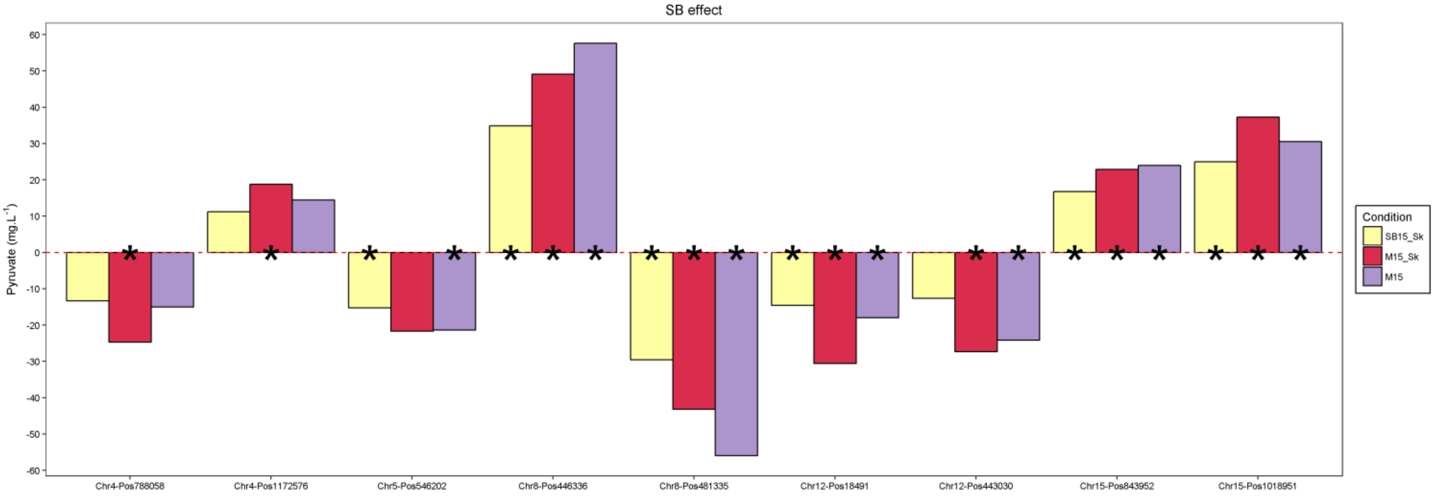

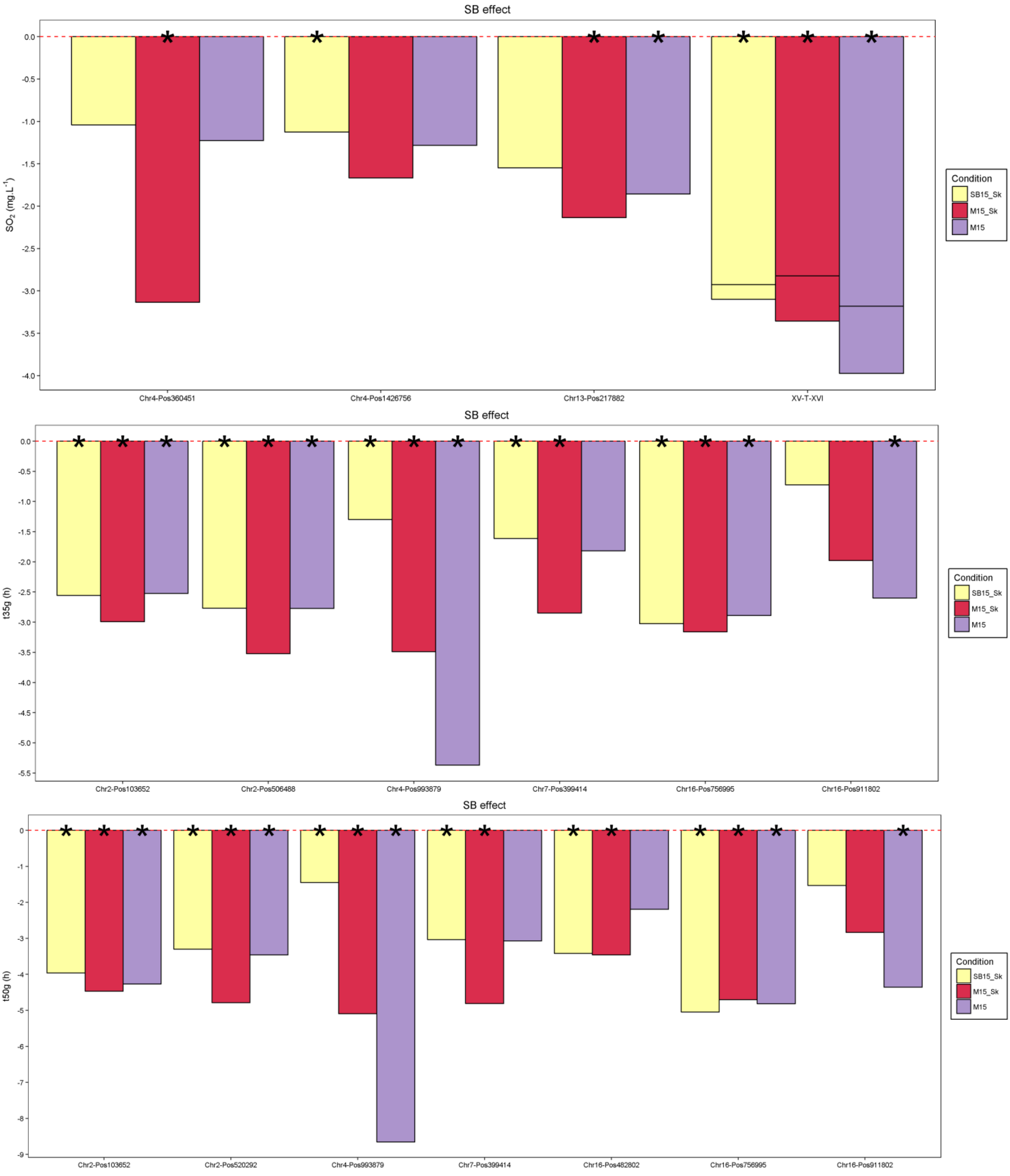

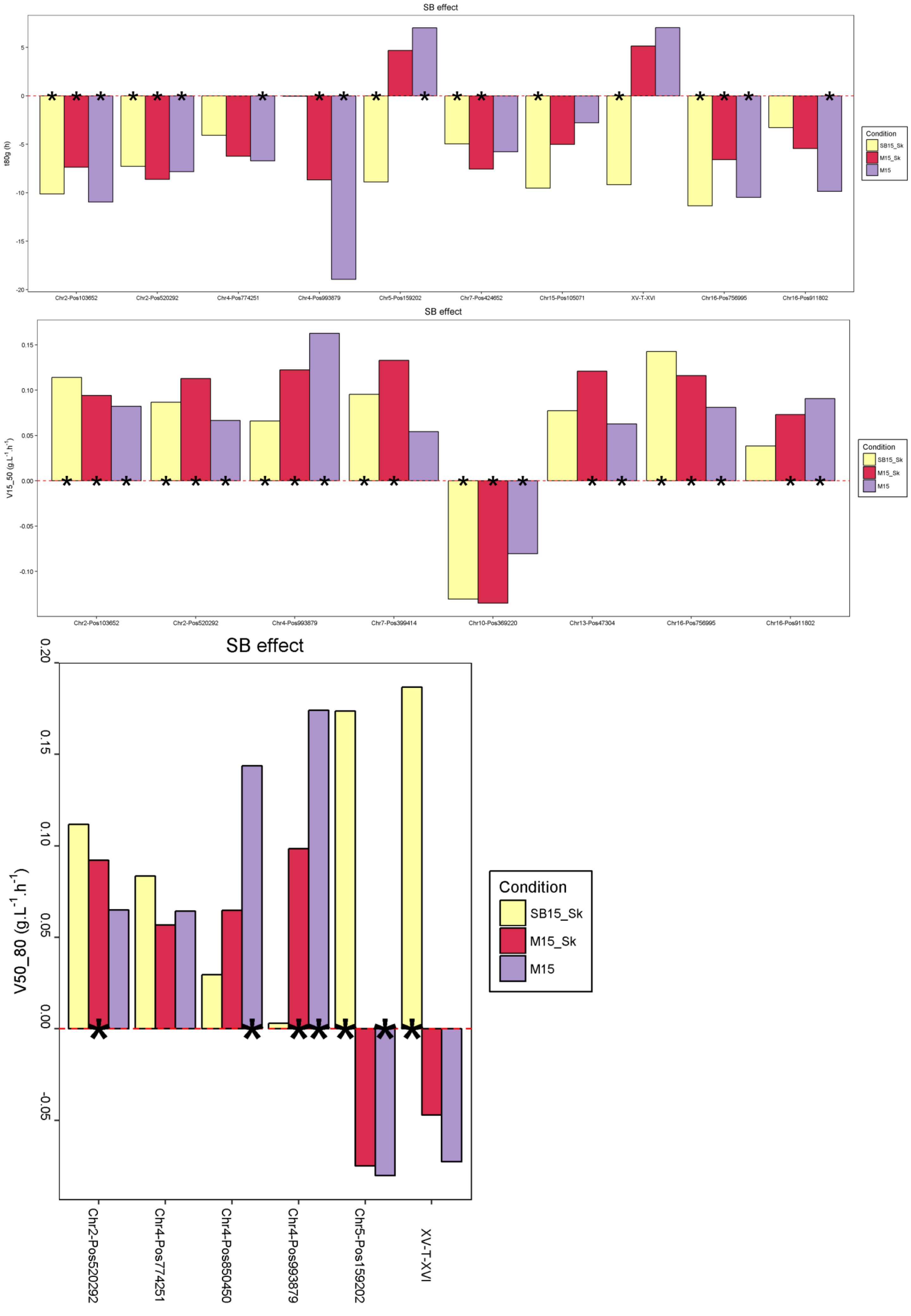

### Supplementary Materials

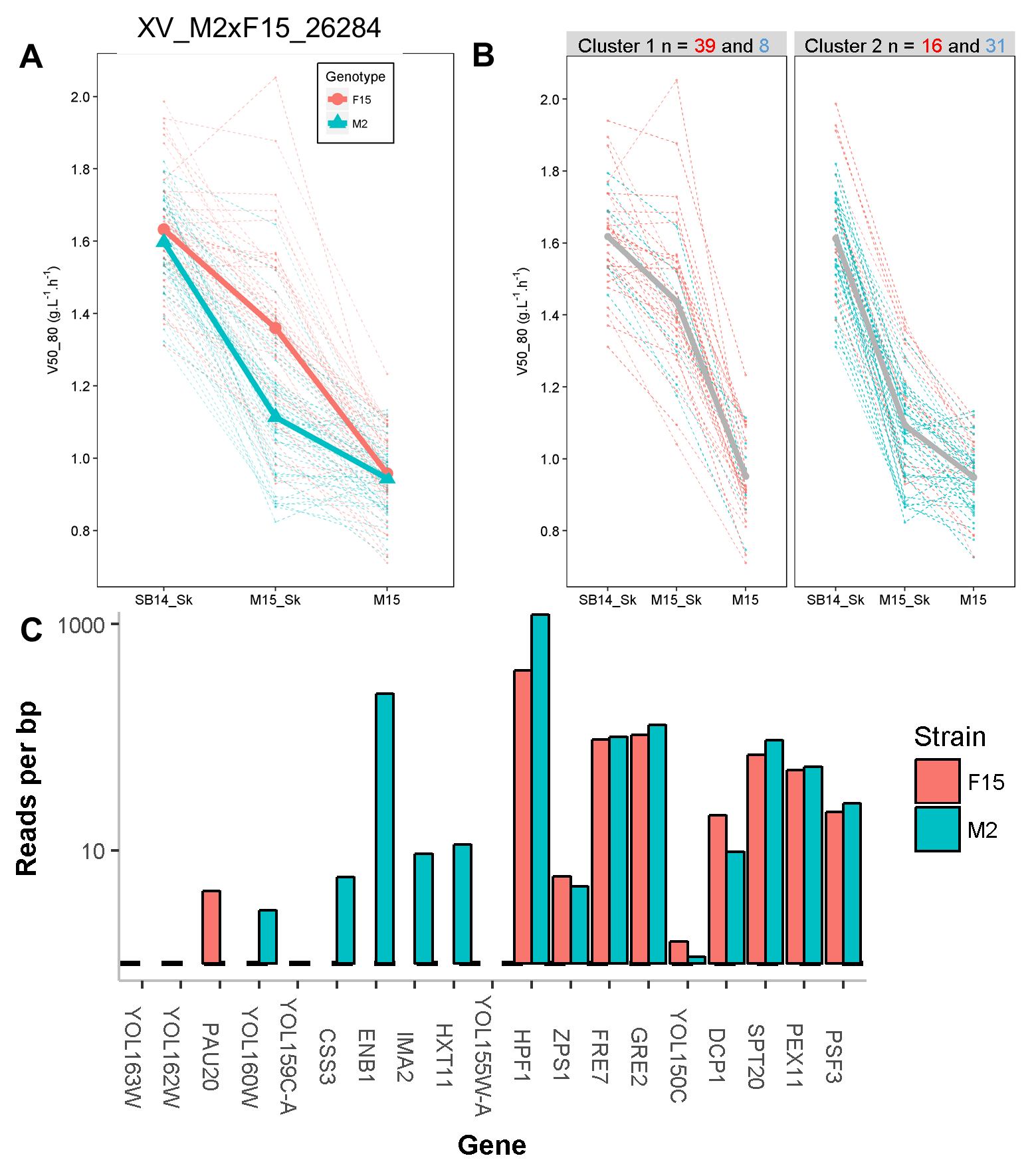

### Supplementary Materials

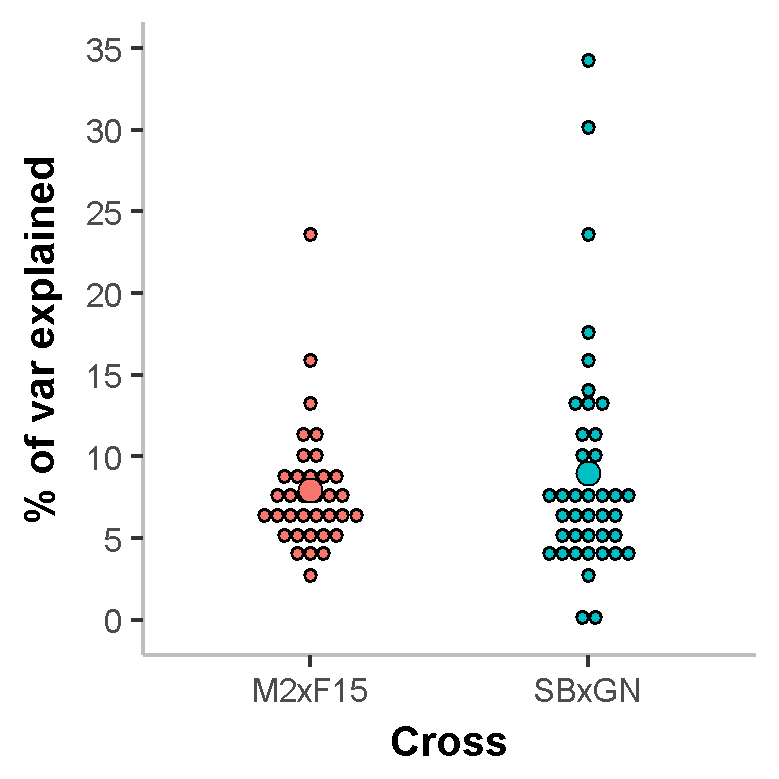
