## Supplementary Materials for "A study of phenotypic plasticity of *Saccharomyces cerevisiae* in natural grape juices shed light on allelic variation under balanced selection"

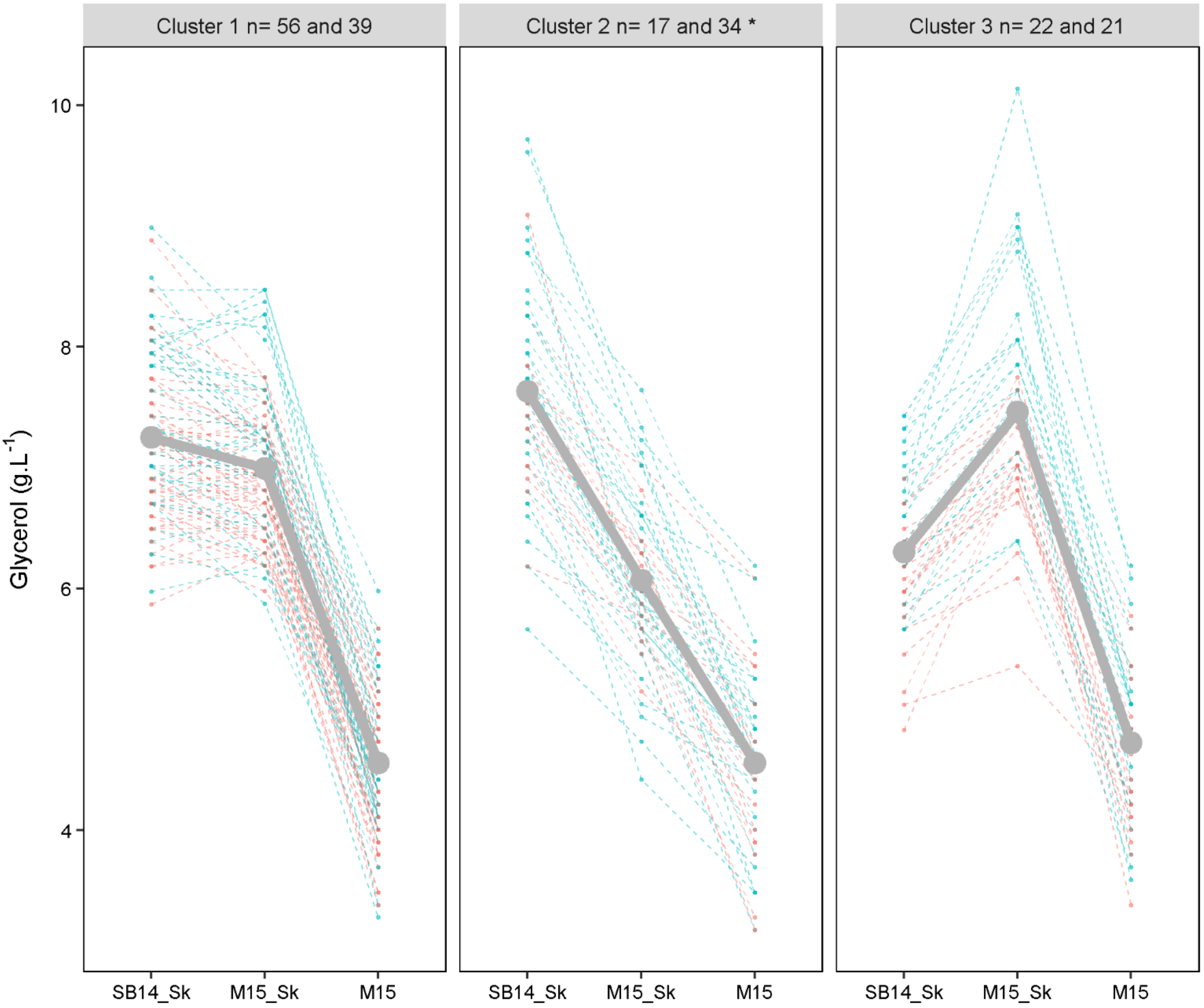
Cluster 1 n= 56 and 39
Cluster 2 n= 17 and 34 *
Cluster 3 n= 22 and 21
Glycerol (g.L$^{-1}$)
10
8
6
4
SB14_Sk
M15_Sk
M15

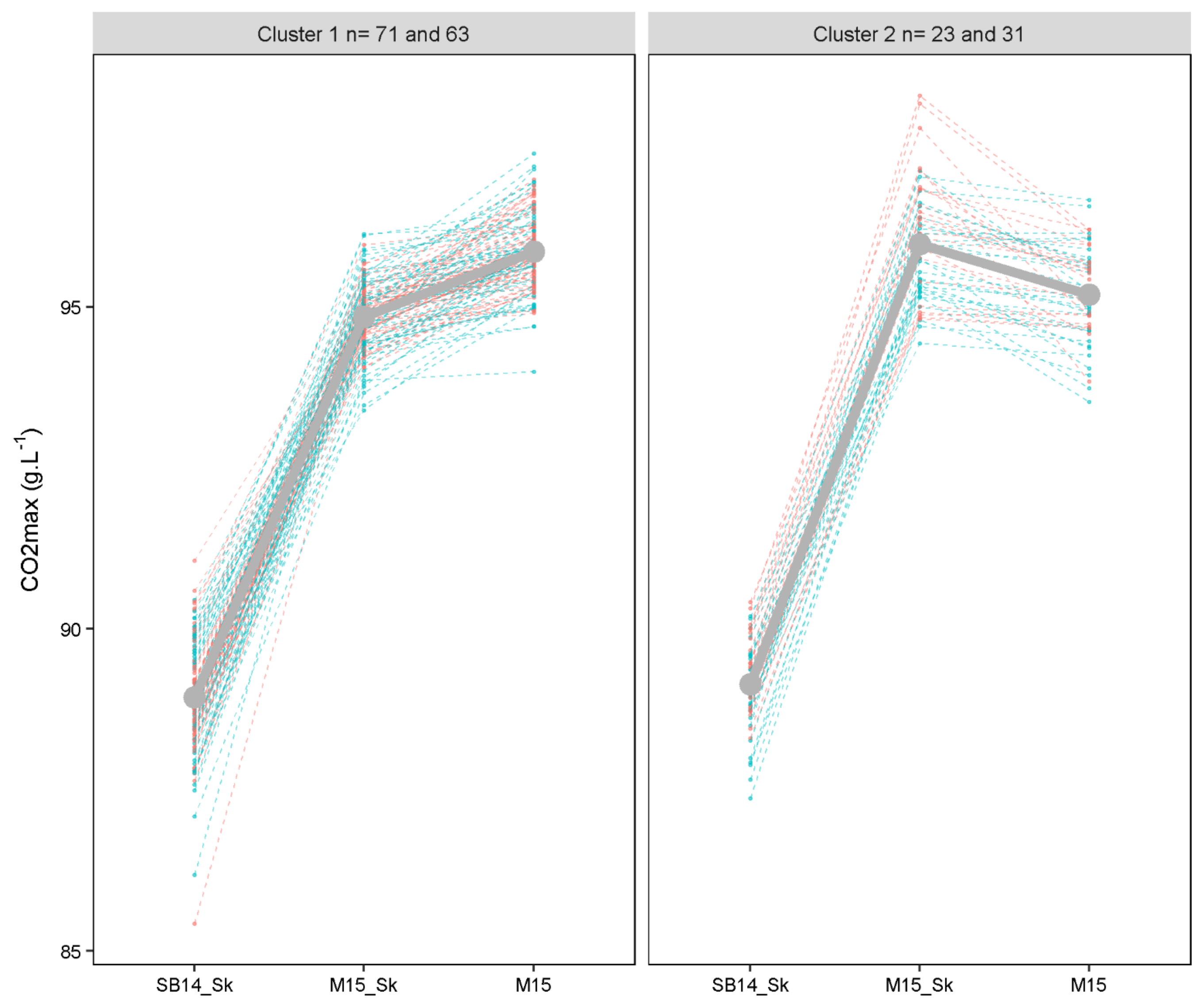

Cluster 1 n= 71 and 63
Cluster 2 n= 23 and 31
CO2max (g.L-1)
95
90
85
SB14_Sk
M15_Sk
M15

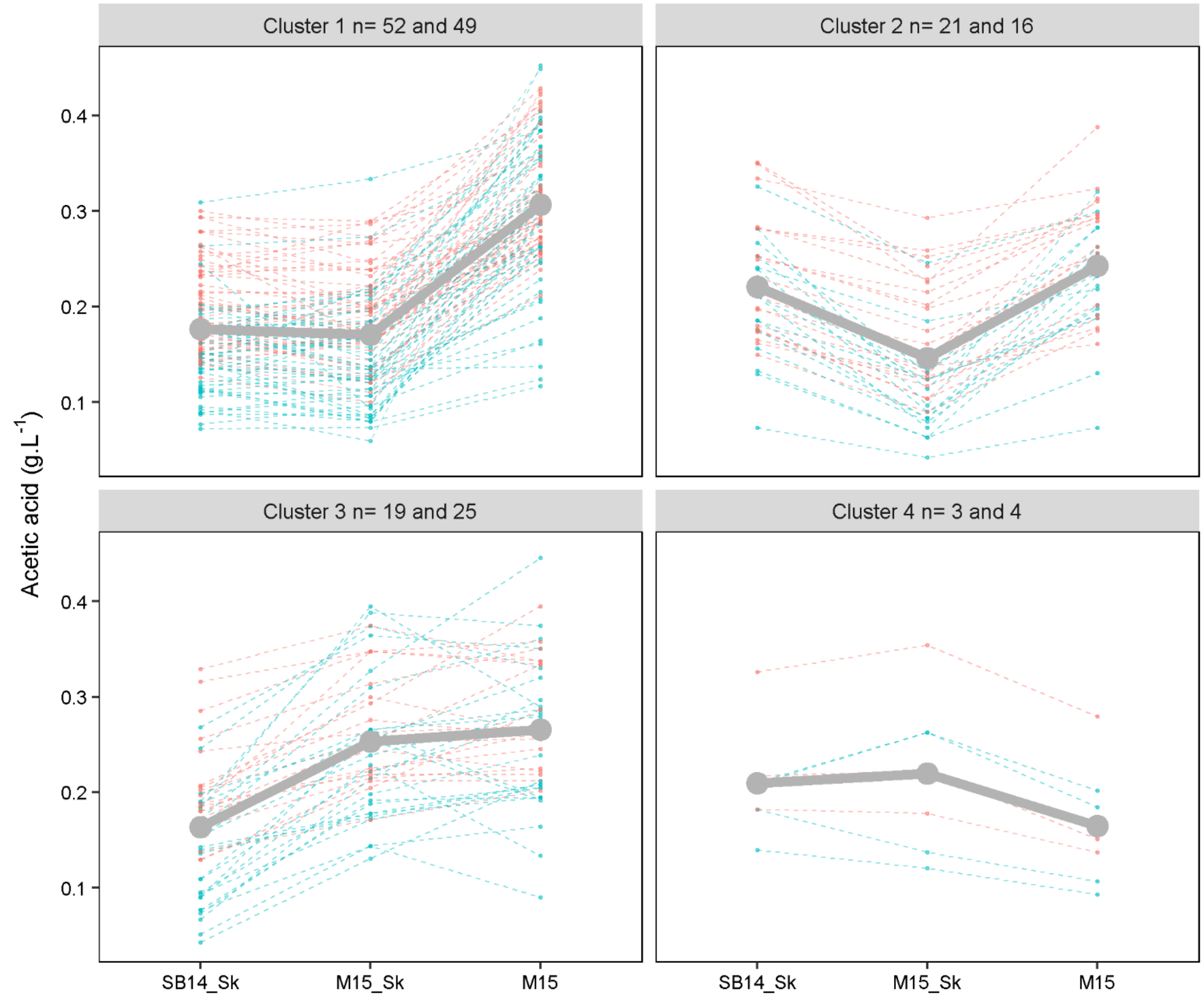

Cluster 1 n= 52 and 49
Cluster 2 n= 21 and 16
Cluster 3 n= 19 and 25
Cluster 4 n= 3 and 4
Acetic acid (g.L-1)
0.1
0.2
0.3
0.4
SB14_Sk
M15_Sk
M15

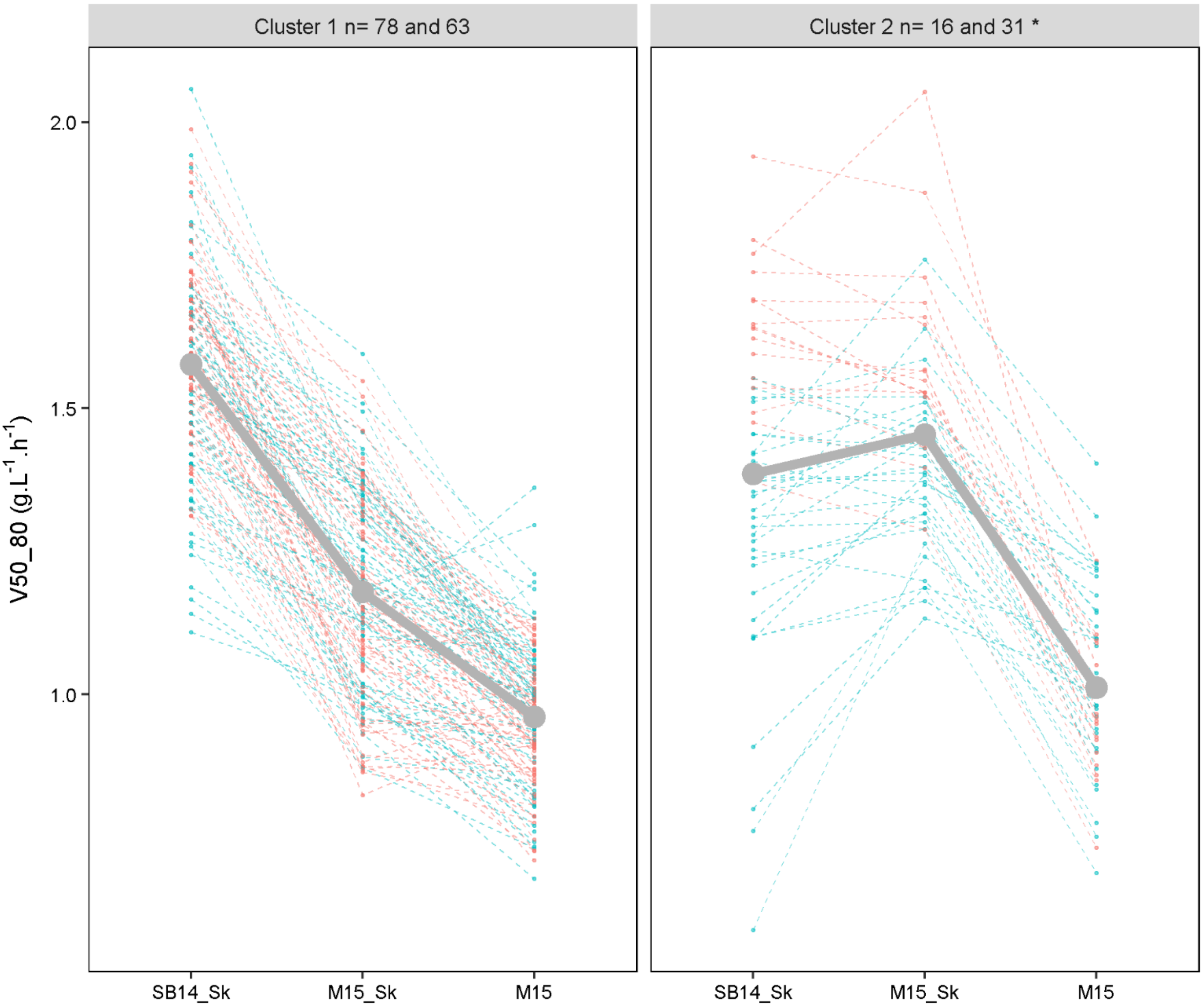

Cluster 1 n= 78 and 63
Cluster 2 n= 16 and 31 *
V50_80 (g.L-1.h-1)
2.0
1.5
1.0
SB14_Sk
M15_Sk
M15
SB14_Sk
M15_Sk
M15

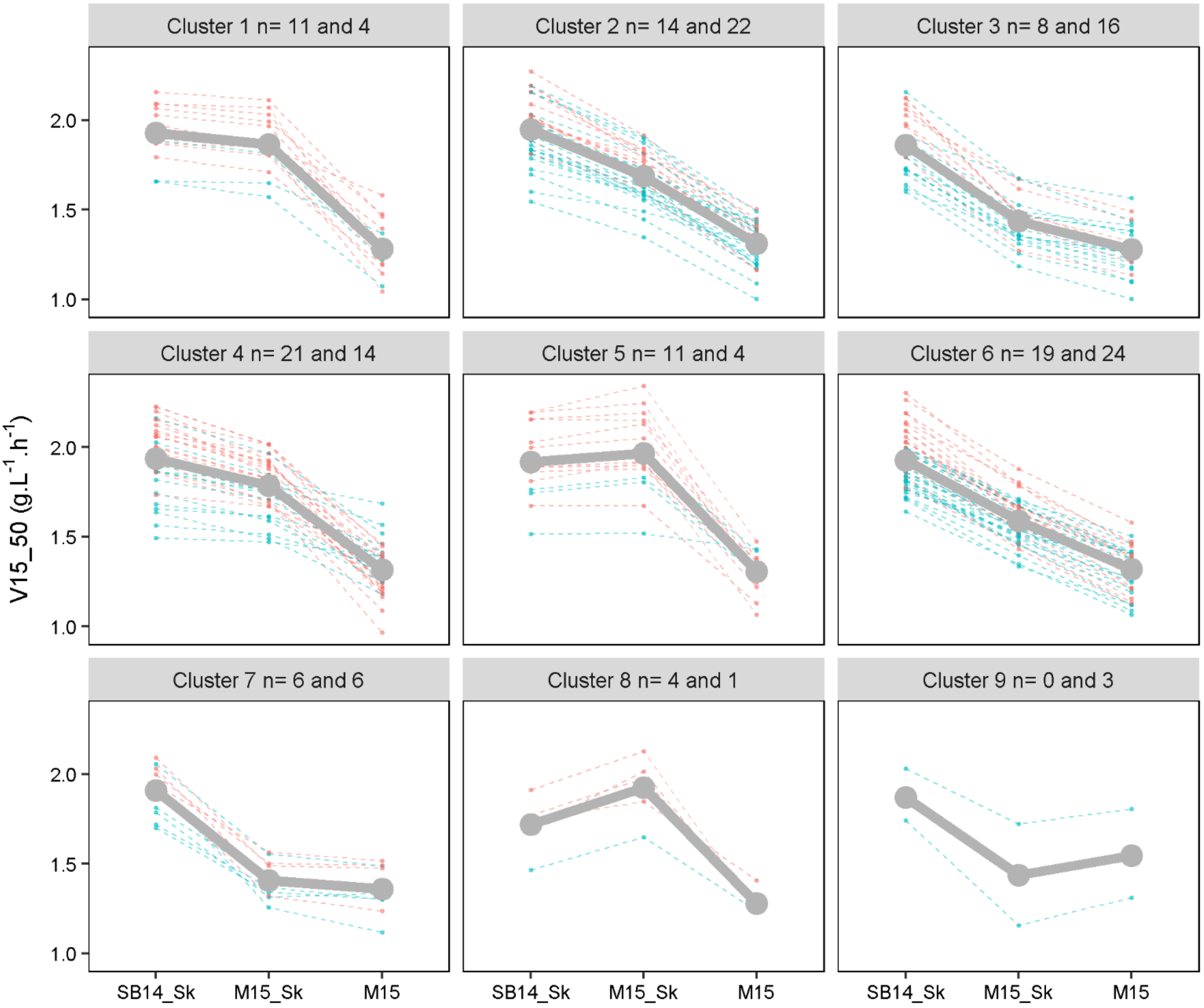

Cluster 1 n= 11 and 4
Cluster 2 n= 14 and 22
Cluster 3 n= 8 and 16
Cluster 4 n= 21 and 14
Cluster 5 n= 11 and 4
Cluster 6 n= 19 and 24
Cluster 7 n= 6 and 6
Cluster 8 n= 4 and 1
Cluster 9 n= 0 and 3
V15_50 (g.L-1.h-1)
1.0
1.5
2.0
SB14_Sk
M15_Sk
M15

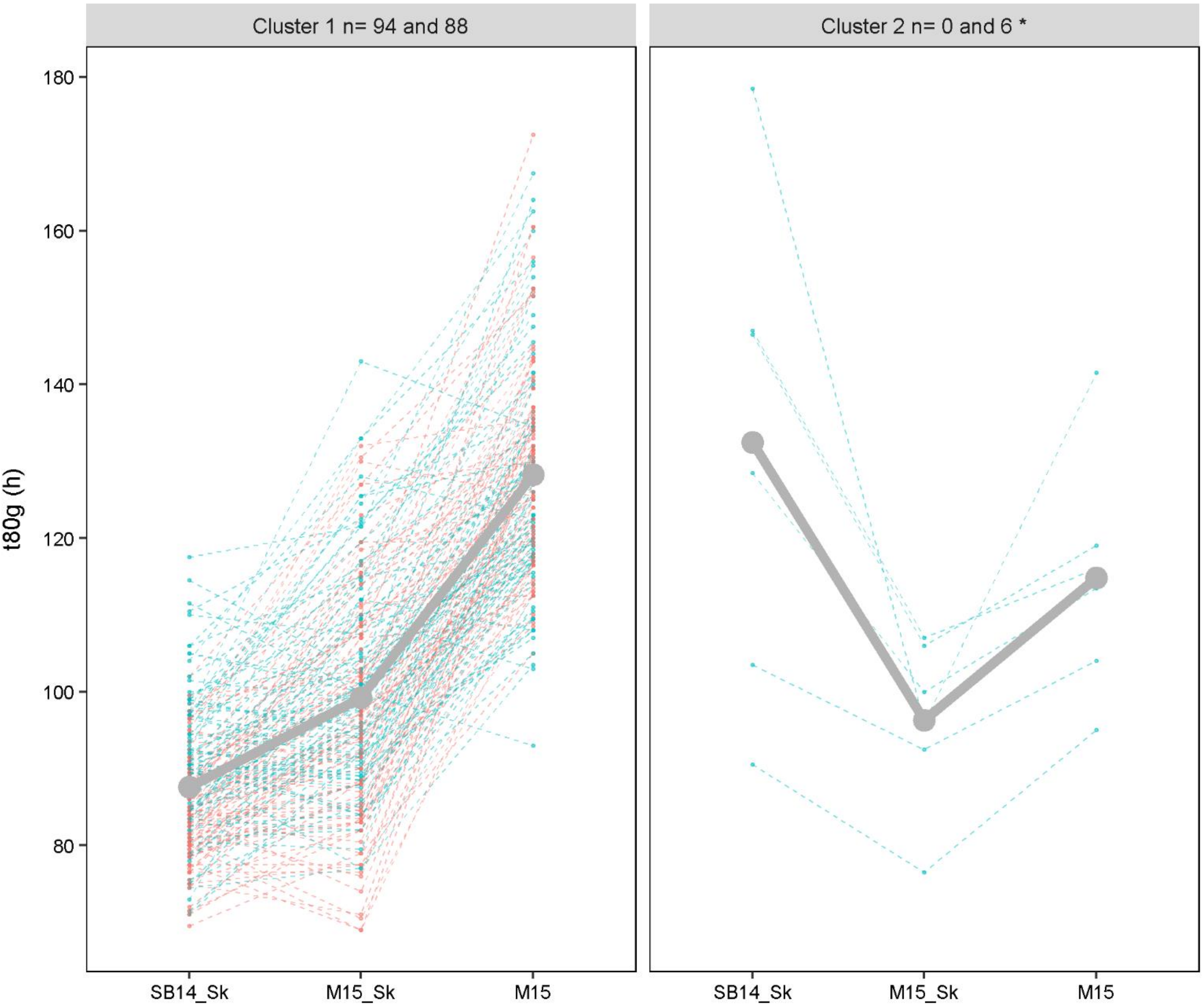

Cluster 1 n= 94 and 88
Cluster 2 n= 0 and 6 *
t80g (h)
180
160
140
120
100
80
SB14_Sk
M15_Sk
M15
SB14_Sk
M15_Sk
M15

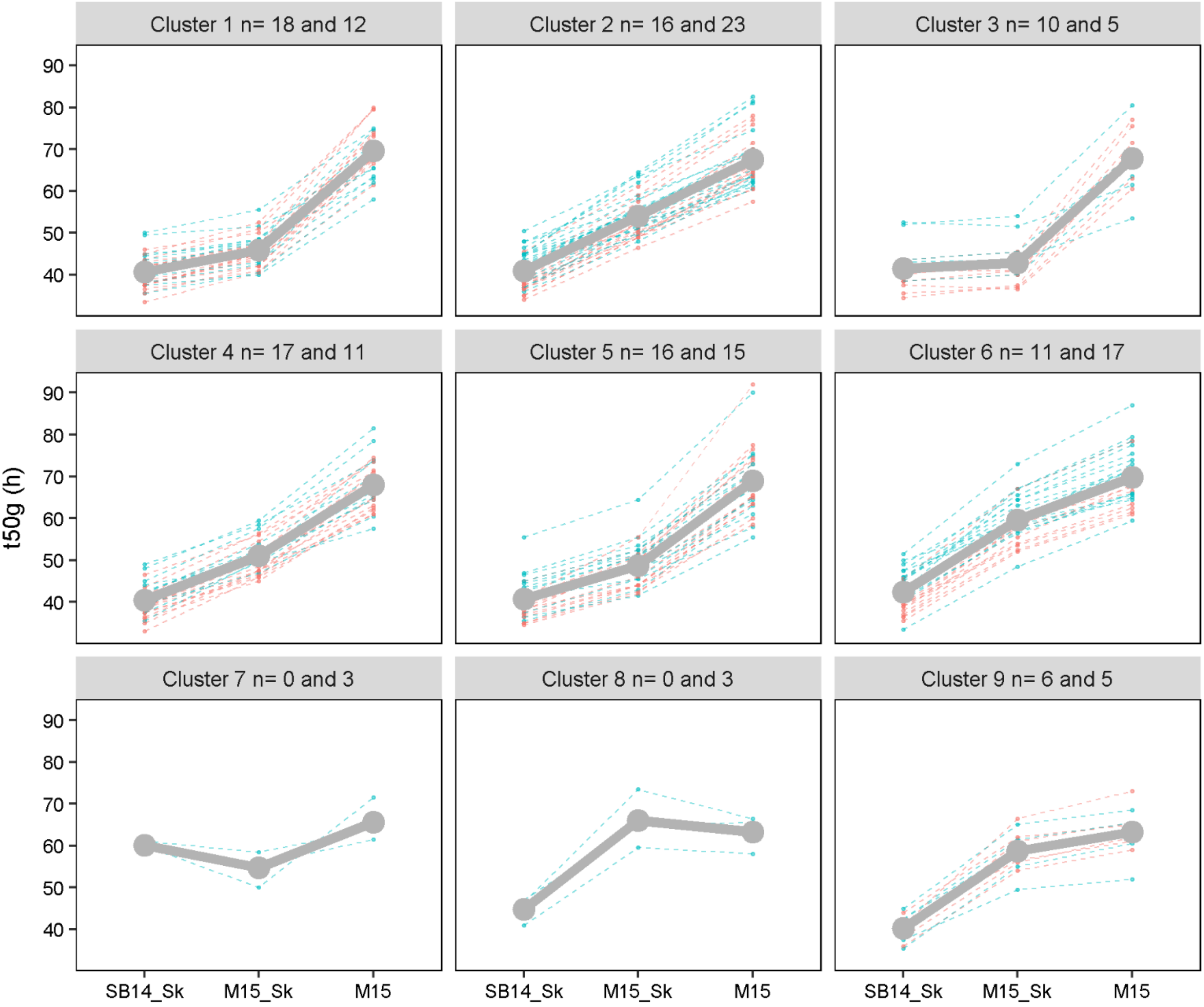
Cluster 1 n= 18 and 12
Cluster 2 n= 16 and 23
Cluster 3 n= 10 and 5
Cluster 4 n= 17 and 11
Cluster 5 n= 16 and 15
Cluster 6 n= 11 and 17
Cluster 7 n= 0 and 3
Cluster 8 n= 0 and 3
Cluster 9 n= 6 and 5
t50g (h)
40
50
60
70
80
90
SB14_Sk
M15_Sk
M15

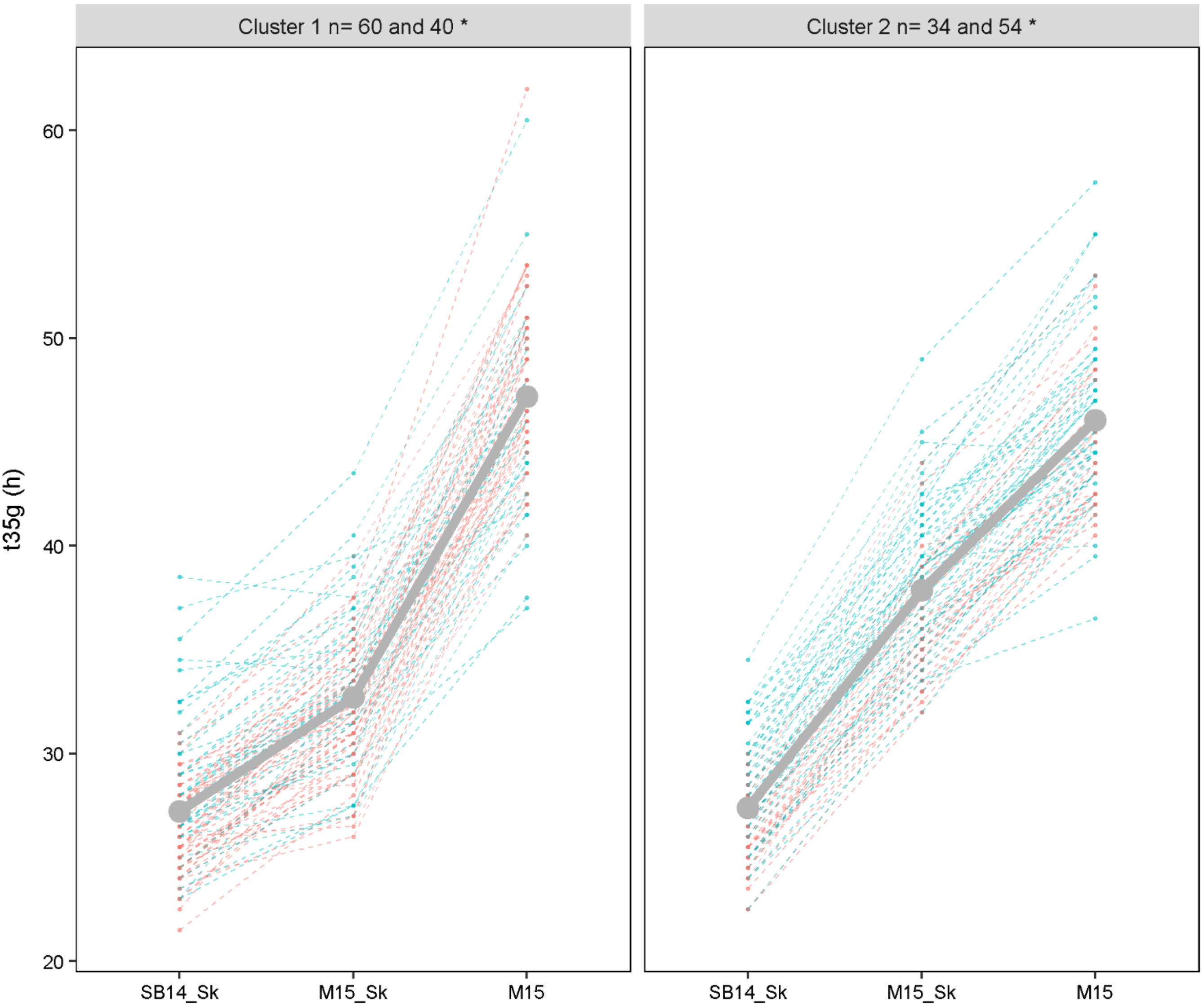
Cluster 1 n= 60 and 40 *
Cluster 2 n= 34 and 54 *
t35g (h)
60
50
40
30
20
SB14_Sk
M15_Sk
M15

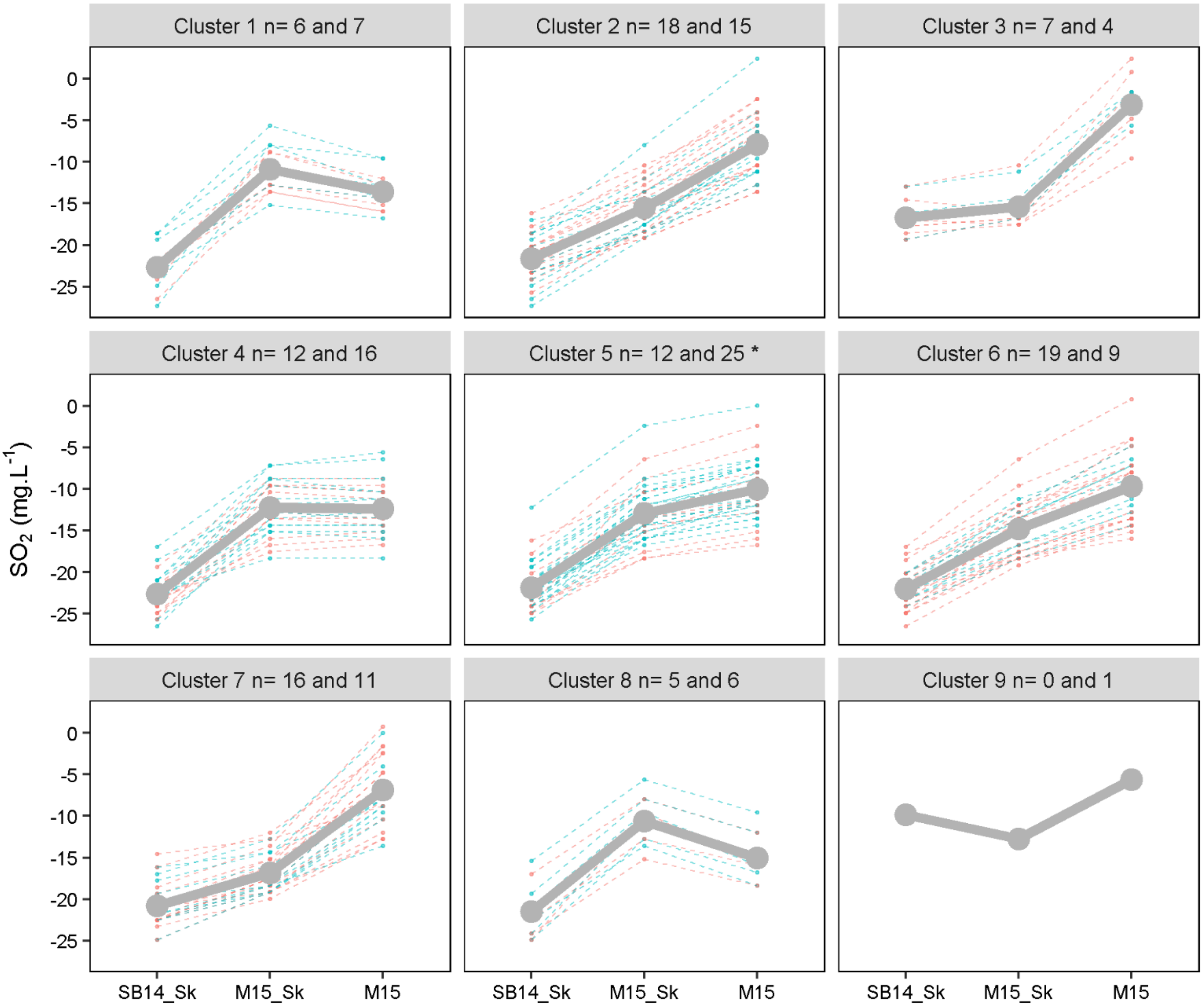

Cluster 1 n= 6 and 7
Cluster 2 n= 18 and 15
Cluster 3 n= 7 and 4
Cluster 4 n= 12 and 16
Cluster 5 n= 12 and 25 *
Cluster 6 n= 19 and 9
Cluster 7 n= 16 and 11
Cluster 8 n= 5 and 6
Cluster 9 n= 0 and 1
$SO_2$ (mg.L$^{-1}$)
0
-5
-10
-15
-20
-25
SB14_Sk
M15_Sk
M15

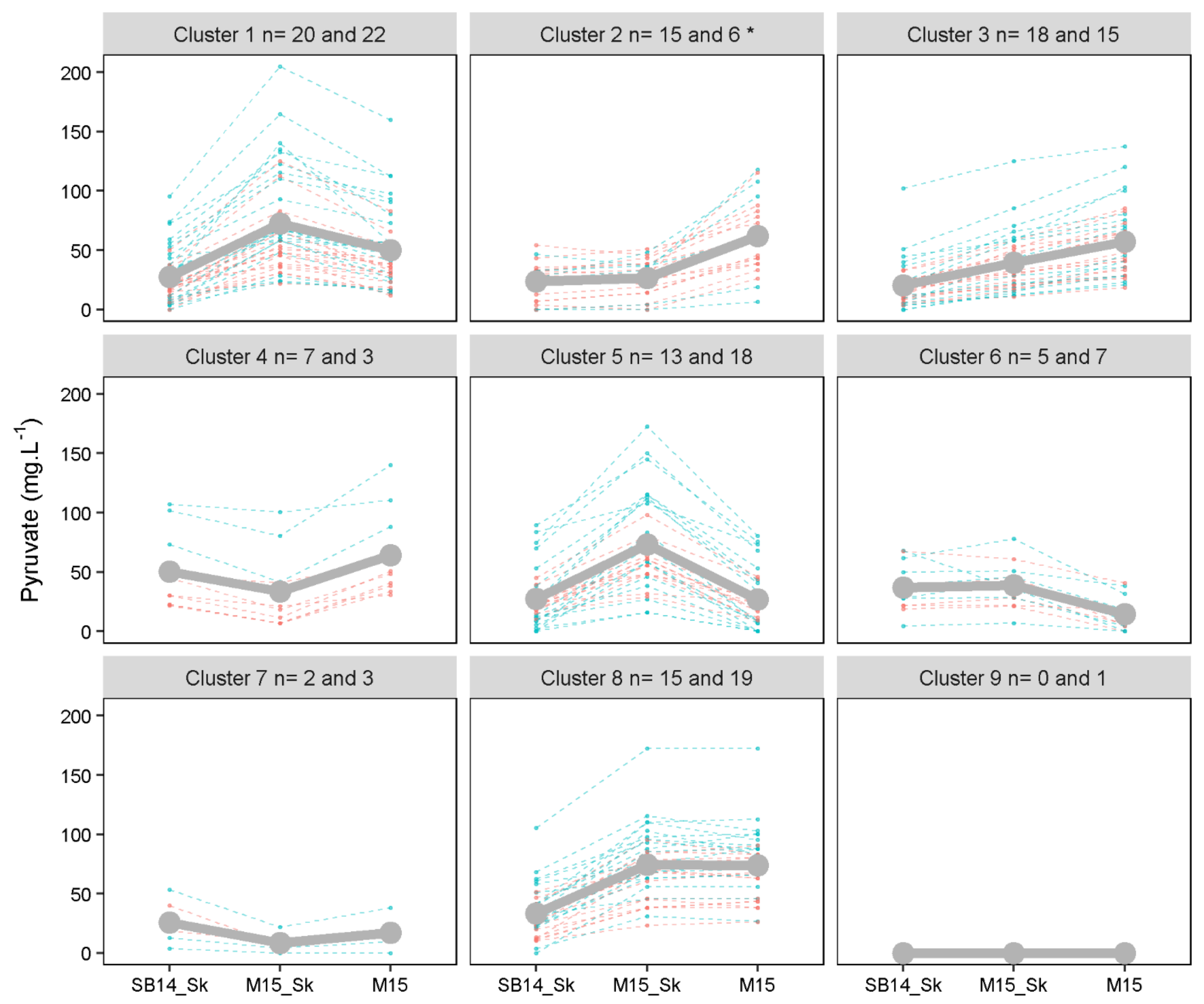

Cluster 1 n= 20 and 22
Cluster 2 n= 15 and 6 *
Cluster 3 n= 18 and 15
Cluster 4 n= 7 and 3
Cluster 5 n= 13 and 18
Cluster 6 n= 5 and 7
Cluster 7 n= 2 and 3
Cluster 8 n= 15 and 19
Cluster 9 n= 0 and 1
Pyruvate (mg.L-1)
0
50
100
150
200
SB14_Sk
M15_Sk
M15

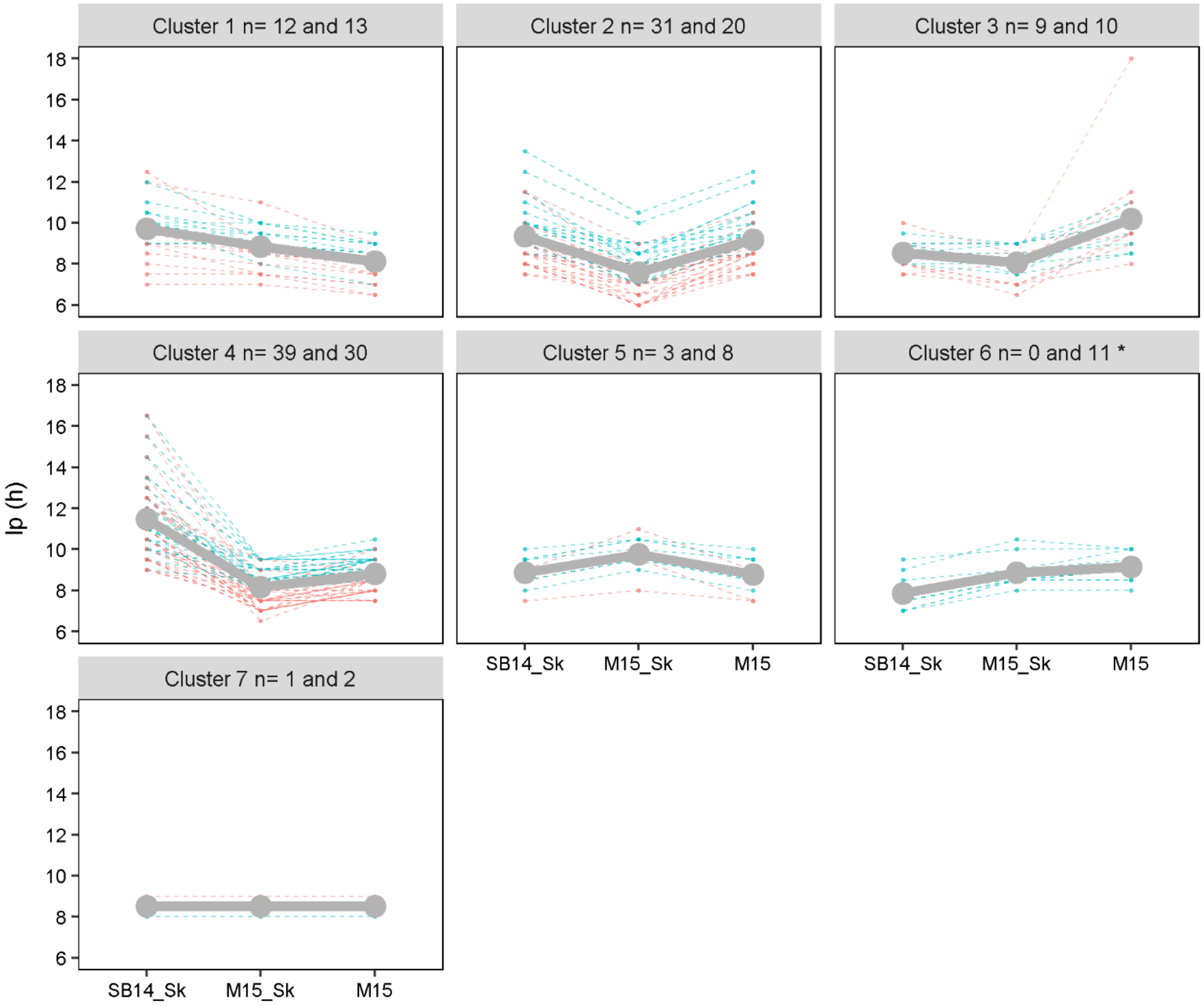
Cluster 1 n= 12 and 13
Cluster 2 n= 31 and 20
Cluster 3 n= 9 and 10
Cluster 4 n= 39 and 30
Cluster 5 n= 3 and 8
Cluster 6 n= 0 and 11 *
Cluster 7 n= 1 and 2
lp (h)
18
16
14
12
10
8
6
SB14_Sk
M15_Sk
M15
